# Blocking a dead-end assembly pathway creates a point of regulation for the DEAD-box ATPase Has1 and prevents platform misassembly

**DOI:** 10.1101/2021.09.06.459192

**Authors:** Xin Liu, Haina Huang, Katrin Karbstein

## Abstract

Assembly of ribosomal subunits occurs via parallel pathways, which accelerate the process and render it more robust. Nonetheless, *in vitro* analyses have also demonstrated that some assembly pathways are dead-ends, presumably due to rRNA misfolding. If and how these non-productive pathways are avoided during assembly *in vivo* remains unknown. Here we use a combination of biochemical, genetic, proteomic and structural analyses to demonstrate a role for assembly factors in biasing the folding landscape away from non-productive intermediates. By binding Rrp36, Rrp5 is prevented from forming a premature interaction with the platform, which leads to a dead-end intermediate, and a misassembled platform that is functionally defective. The DEAD-box ATPase Has1 separates Rrp5 and Rrp36, allowing Rrp5 to reposition to the platform, thereby promoting ribosome assembly and enabling rRNA processing. Thus, Rrp36 establishes an ATP-dependent regulatory point that ensures correct platform assembly by opening a new folding channel that avoids funnels to misfolding.

## (Introduction)

Ribosomes are highly intricate and universally conserved RNA-protein complexes that catalyze protein synthesis in all cells. Made up of 4 RNAs and 79 ribosomal proteins in eukaryotes, their assembly proceeds via multiple assembly routes(Cheng et al., 2019; Cheng et al., 2020; Davis et al., 2016; Du et al., 2020; Mulder et al., 2010; Sanghai et al., 2018; Sashital et al., 2014). How these routes are populated to avoid non-productive pathways that involve kinetically trapped misfolded assembly intermediates, a problem common in RNA folding(Ganser et al., 2019; Herschlag et al., 2018; Woodson, 2011), remains unclear, as is the role that the roughly 200 assembly factors (AFs) that are required for ribosome assembly play in this process. This problem is exacerbated because our knowledge about the structure of misfolded intermediates and thus the causes for misfolding of large RNAs such as rRNAs remains limited.

While many structures of ribosome assembly intermediates are now available, they do not typically reveal misassembled or dead-end intermediates, likely because these are avoided within cells and/or rapidly degraded. One exception is observed on the platform of the assembling small subunit, which contains the mRNA exit channel and the tRNA exit (E-site). The available structures show two classes of structures that differ by the already formed RNA elements and the bound ribosomal proteins (**Figures 1A&B**, (Barandun et al., 2017; Cheng *et al*., 2019; Cheng et al., 2017; Cheng *et al*., 2020; Du *et al*., 2020; Sun et al., 2017)). The class that can mature further (top in **Figures 1A&B**, (Cheng *et al*., 2019; Du *et al*., 2020), and is the dominant class in intermediates grown in rich media, lacks helix 24 (h24) but contains the ribosomal proteins Rps1, Rps27 and Rps22/uS8. The other subclass (bottom in **Figures 1A&B)**, which is non-productive and the only class found when intermediates are isolated from cells in stationary phase, lacks these two ribosomal proteins. In contrast, h24 is prematurely formed and misoriented, due to its binding to Rrp5, thereby blocking further platform maturation. How this unproductive pathway, characterized by premature binding of Rrp5 to the platform, is largely avoided in well-fed yeast cells remains unknown.

**Figure 1.**
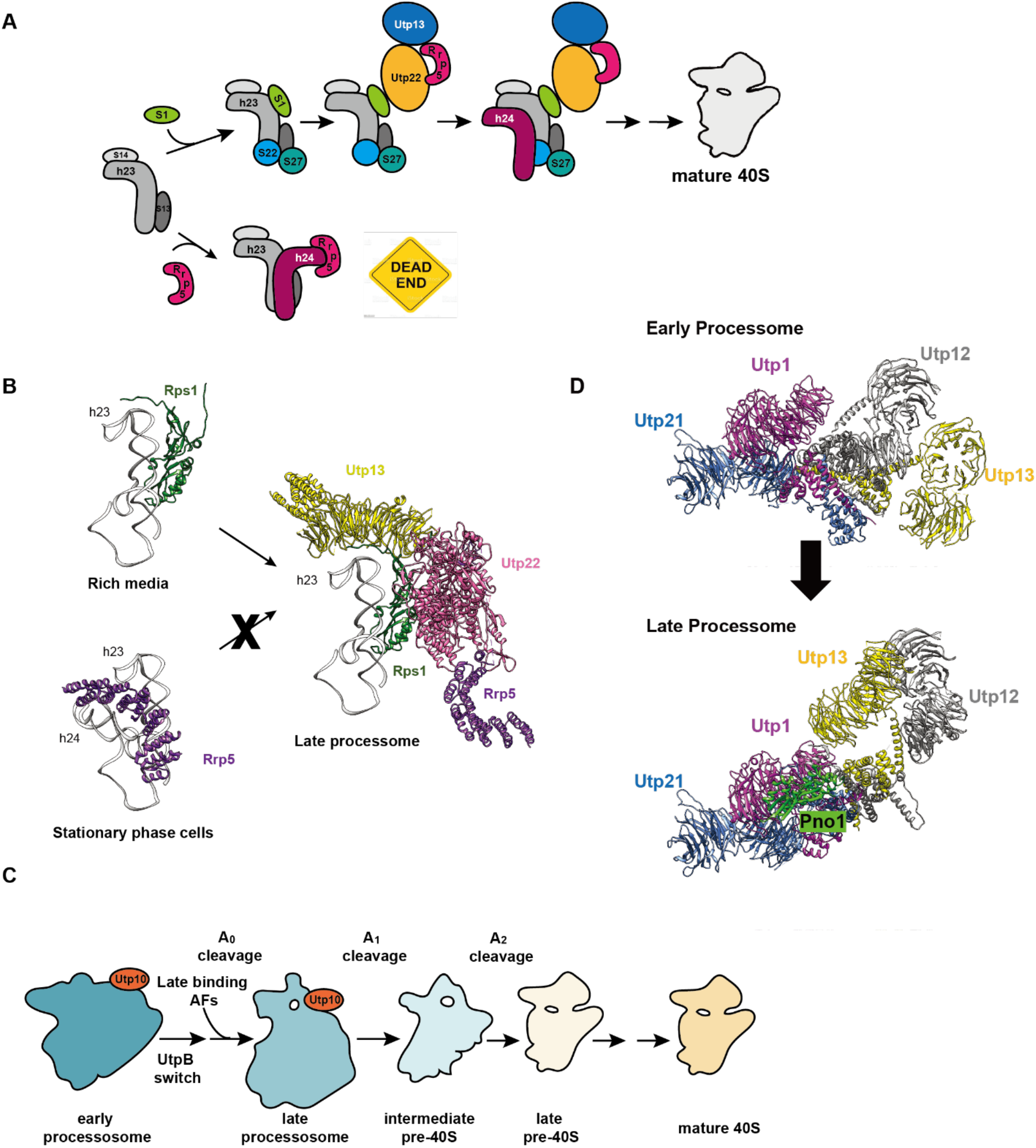
rRNA processing is linked to compaction of early 40S assembly intermediates and the UtpB conformational switch. A. Two pathways for assembly of the platform are visualized in cryo-EM structures. (Top) in cells grown in rich media Rps1, Rps27 and Rps22/uS8 bind the platform helix 23, which already is bound to Rps13/uS15 and Rps14/uS11. After the UtpB switch, Utp13 binds Utp22, stabilized by Rrp5. Eventually h24 forms and maturation proceeds. Based on PDB ID 6ND4, 6LQQ, 6LQP, 6LQS; (Bottom) in stationary phase cells, Rrp5 stabilizes h24 prematurely, by misorienting it. These intermediates cannot mature. Based on PDB ID 5WLC. B. Rrp5 has two positions on the platform: the productive position near Utp13 (PDB ID 6LQQ) and a non-productive position (PDB ID 5WLC) that misorients h24. C. Assembly of the processosome is integrated with rRNA processing. Early processosomes convert to late processosomes after transcription of the 3’-end of 18S rRNA, and a switch in UtpB, which enables the recruitment of a set of late-binding assembly factors (Chaker-Margot *et al*., 2015; Chen *et al*., 2020; Hunziker *et al*., 2019; Zhang *et al*., 2016). This allows processing at site A_0_. Subsequent processing at the A_1_ and A_2_ sites is accompanied by additional changes in processosome structure and composition(Cheng *et al*., 2020; Du *et al*., 2020). D. In the conversion from early to late 90S precursors, UtpB undergoes a conformational switch in Utp1/Utp21, which repositions Utp12 and Utp13 relative to the rest of UtpB and the rest of the processosome. This creates the Pno1 binding site (Hunziker *et al*., 2019). Figures were made from PDB ID 6ND4 and 6LQP.

Along the productive pathway, structural studies have revealed multiple conformational transitions as the subunit assembles from an early processosome via a late processosome, into intermediate and then late pre-40S intermediates (**Figure 1C**). These transitions are integrated with rRNA processing steps (**Figure 1C, S1A**). Thus, the early processosome is blocked prior to the first rRNA cleavage step at so-called site A_0_ (Hunziker et al., 2019), while the next intermediate, the late processosome, is processed at the A_0_ site, but stalled prior to the next step, cleavage at site A_1_ (Barandun *et al*., 2017; Cheng *et al*., 2020; Du *et al*., 2020). Conversion of the early to the late processosome is linked not just to A_0_ processing, but also to a switch in the UtpB subcomplex (composed of Utp1, Utp21, Utp12, Utp13 and Utp6 and Utp18): by changing the Utp1 and Utp21 interface, their interaction partners Utp12 and Utp13 are rotated to a vastly different location with respect to the rest of UtpB and the processosome **(Figure 1D**, (Hunziker *et al*., 2019)). This enables the recruitment of a subset of late-binding processosome factors, which ultimately enable rRNA processing (Chaker-Margot et al., 2015; Chen et al., 2020; Hunziker *et al*., 2019; Zhang et al., 2016). How this transition is promoted in cells remains entirely unknown.

DEAD-box proteins are RNA-dependent ATPases (**Figure S1B**, (Cordin et al., 2006; Jarmoskaite and Russell, 2014; Linder and Jankowsky, 2011)). While commonly referred to as RNA helicases, it is now clear that most, if not all, lack processive unwinding activity, but instead release RNA-binding proteins, locally unfold helices, or even catalyze duplex annealing (Mallam et al., 2012; Putnam and Jankowsky, 2013; Young et al., 2013). While DEAD-box proteins are ubiquitous regulators of RNA-dependent biological processes, the largest subgroup of them is involved in ribosome assembly (Martin et al., 2013). What roles they play in this process remains largely unknown, despite their obvious candidacy for regulating conformational transitions by releasing assembly factors or changing RNA structure.

To understand how premature binding of Rrp5 at the platform was avoided in rich media to promote productive assembly, we used a combination of biochemical, genetic, mass spectrometry and structure mapping experiments. These experiments show that in very early assembly intermediates Rrp5 is tethered away from the platform via an interaction with the assembly factor Rrp36 on the nascent body. This sets up a point of regulation for the DEAD-box ATPase Has1, which uses ATP to release Rrp36 from nascent subunits, thereby enabling the repositioning of Rrp5 from the body to the platform. Because the interactions of Rrp5 at the platform stabilize the UtpB switch, the Has1-mediated repositioning of Rrp5 enables the UtpB switch, allowing for the recruitment of late-binding processosome factors, and thereby the activation of rRNA processing. Thus, Rrp5 and Rrp36 cooperate to establish a checkpoint that is regulated in an ATPase-dependent manner by the DEAD-box protein Has1 to ensure proper maturation of the platform. Accordingly, disrupting this checkpoint leads to platform misassembly. This work reveals a novel function for assembly factors in blocking non-productive assembly channels.

## Results

### Rrp5 binds Rrp36 via its N-terminal S1 domain

Previous biochemical and yeast two hybrid analyses indicated an interaction between Rrp5 and the Rrp9/Rrp36 complex located at the nascent body (Clerget et al., 2020). This puts Rrp5 ∼140Å away from the better-characterized binding site on the platform, where cryo-EM structures have located Rrp5 and its binding partner Utp22(Barandun *et al*., 2017; Du *et al*., 2020), and where we show it interacts with Krr1 (**Figure S2**). Moreover, a careful comparison of these cryo-EM structures indicated two distinct positions of Rrp5 on the platform (**Figure 1A&B**): either the Rrp5 tetratricopeptide repeat (TPR) domain (see **Figure S1C** for a schematic of Rrp5’s domain organization) was bound to h24, holding it in the wrong orientation and thus leading to a dead-end assembly intermediate ((Barandun *et al*., 2017; Du *et al*., 2020; Sun *et al*., 2017). Alternatively, the TPR domain was bound to Utp22 and Utp13 (Cheng *et al*., 2019; Cheng *et al*., 2017; Cheng *et al*., 2020; Du *et al*., 2020; Sun *et al*., 2017). In these structures, platform assembly had progressed further with the incorporation of Rps1, as well as Rps22/uS8 and Rps27 (**Figure 1A&B**, both classes contain Rps13(uS15) and Rps14 (uS11)). We thus hypothesized that the binding of Rrp5 to Rrp36/Rrp9 on the body served to delay binding of Rrp5 to the platform, thereby funneling 40S assembly intermediates into a productive assembly pathway.

To test this model, we first sought to better characterize the interaction between Rrp5 and Rrp36/Rrp9 on the assembling body. Yeast two-hybrid and protein interaction assays had shown that Rrp36 bound directly to Rrp5, as well as to Rrp9 (Clerget *et al*., 2020). Thus, we first determined which of the subdomains in Rrp5 was responsible for binding to Rrp36. For these experiments we expressed and purified recombinant Rrp5 and Rrp36 from *E. coli*, and then used pulldown assays to confirm their binding. As expected, these experiments demonstrate a strong interaction between Rrp5 and Rrp36 (**Figure 2A**). Next, we tested different fragments of Rrp5 for their ability to bind Rrp36. Removal of either the first two, or just the first S1 domain abolished the interaction with Rrp36 (**Figure 2B, Figure S3A)**, suggesting strongly that the first S1 domain of Rrp5 was responsible for the binding to Rrp36. To confirm this conclusion, we expressed and purified just the first S1 domain alone, or in combination with the second or second and third S1 domains and tested them for binding to Rrp36. These data show that the first S1 domain alone can produce a stoichiometric Rrp5-Rrp36 complex (**Figure 2C**). Together, these data demonstrate that the first S1 domain in Rrp5 is responsible for locating Rrp5 to the Rrp36/Rrp9 complex on the assembling body.

**Figure 2.**
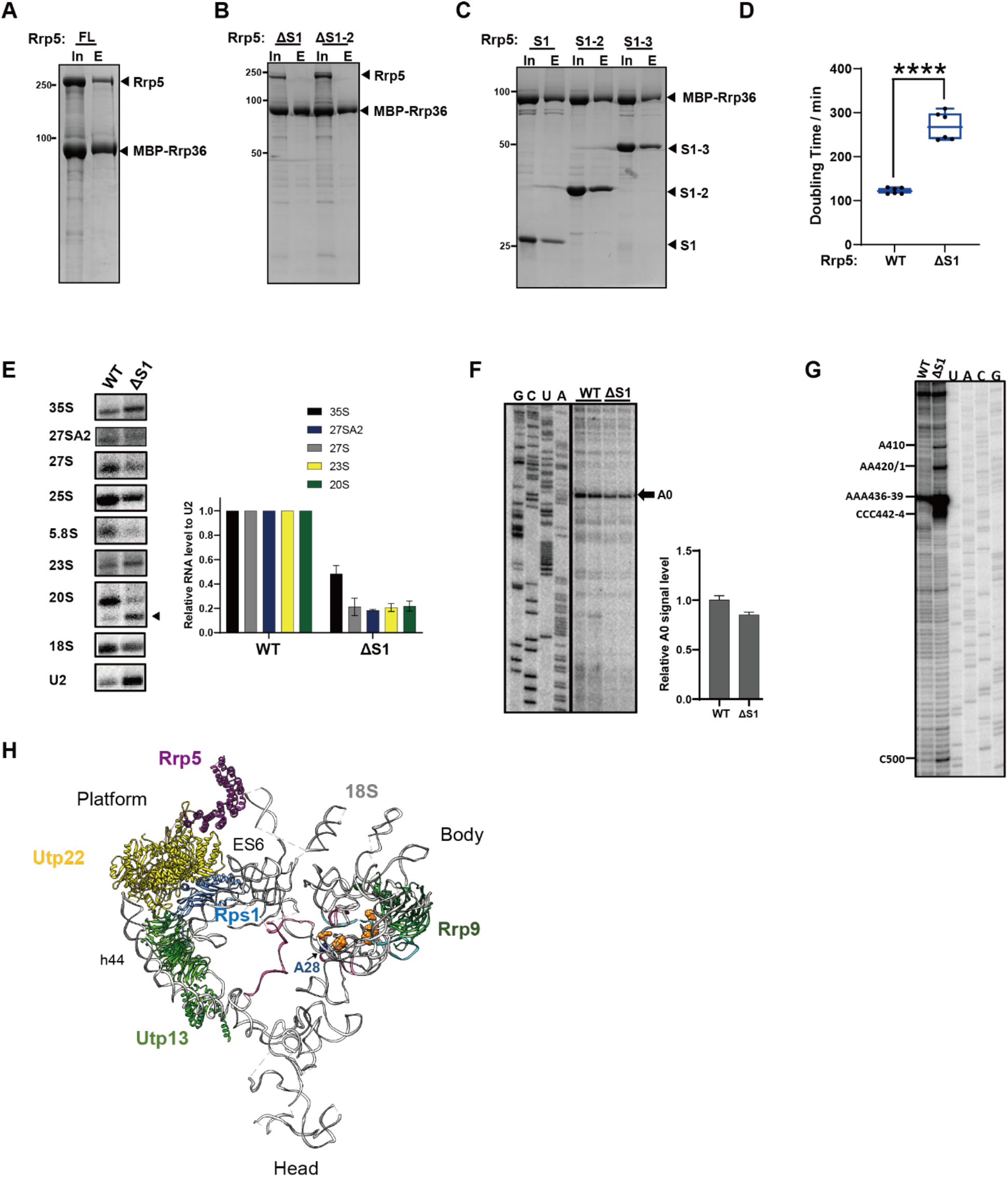
Disrupting the binding of Rrp5 and Rrp36 early in assembly leads to defective body assembly and 40S processing defects. A. Recombinant MBP-Rrp36 immobilized on amylose resin interacts with recombinant purified full length Rrp5 (Rrp5_FL). In: Input; E: Elution. B. Deletion of the first S1 domain in Rrp5, either alone (Rrp5_ΔS1) or in combination with the second one (Rrp5_ΔS1-2), disrupts the interaction with Rrp36. C. The first S1 domain in Rrp5 alone (Rrp5_S1) or in combination with the second (Rrp5_S1-2) and third one (Rrp5_S1-3) binds Rrp36. None of the proteins in A-C interact with amylose-resin (**Figure S3A**). The position of MBP-Rrp36 is indicated with an arrow. D. Doubling times of Gal:Rrp5 cells supplied with plasmids encoding Rrp5 WT or Rrp5_ΔS1. Significance was tested using an unpaired t-test. ****, p<0.0001. n≥6. E. Northern blot analysis of rRNA processing intermediates from Gal:Rrp5 cells expressing plasmid-encoded Rrp5 WT or Rrp5_ΔS1 as in D. The locations of Northern probes are indicated with black bars in **Figure S1A**. U2 serves as loading control. A degradation or mis-cleavage product in Rrp5_ΔS1 is indicated with arrow on the right. For the quantifications rRNA levels were normalized to U2 and to the sample with wild type Rrp5. The data are averages from 3 biological replicates and error bars show standard deviation of the mean. F. Analysis of A_0_ cleavage by reverse transcription of total RNA extracted from Gal:Rrp5-depleted cells expressing plasmid-encoded protein as in D. Shown below is the quantification of A_0_ cleavage band, normalized to the full extension band (not shown). The data are averages from 2 biological and 2 technical replicates and error bars show standard deviation of the mean. G. Reverse transcription maps the 5’-end of the smaller rRNA piece observed in Rrp5_ΔS1 cells in panel E (20S). H. The miscleaved 5’-ends observed in Rrp5_ΔS1 cells are mapped onto the late processosome structure with orange spacefill (PDB ID 6LQQ). A28 is shown in blue spacefill and indicated by arrow. Select assembly factors are shown. Crosslinking sites for Rrp5 are shown in pink.

### Deletion of the first S1-domain in Rrp5 leads to defects in the assembling 40S body

To dissect the importance of locating Rrp5 to the assembling body, we deleted the first S1 domain and then tested its effects on ribosome maturation. For these experiments we used a previously described galactose-inducible Rrp5 strain(Khoshnevis et al., 2016; Khoshnevis et al., 2019; Young and Karbstein, 2011) and supplemented it with a plasmid encoding a fragment of Rrp5, lacking the first S1 domain (Rrp5_ΔS1), and tested its effects on yeast growth (**Figure 2D**). The data demonstrate that the Rrp5_ΔS1 truncation displays a significant growth defect, suggesting an important role for this domain in ribosome assembly.

Because cursory previous analyses had suggested that the Rrp5 N-terminus was required for 60S assembly(Eppens et al., 1999; Lebaron et al., 2013; Torchet et al., 1998; Young and Karbstein, 2011), we first used Northern blot analysis to confirm that the Rrp5_ΔS1 mutant affects 40S processing. Indeed, deletion of the first S1 domain in Rrp5 leads to strong depletion of the A_2_ cleavage products, 20S rRNA and 27S rRNA (**Figure 2E**, see **Figure S1A** for a scheme describing the rRNA processing pathway), demonstrating a role for this element in early 40S processing. Primer extension indicates that A_0_ cleavage is largely unaffected in this mutant (**Figure 2F**). Moreover, we observe the accumulation of a smaller product below 20S rRNA. This fragment is detected by probes in ITS1, the segment of precursor RNA between 18S and 5.8S rRNAs (**Figure S1A**), as well as within the 5’-end of 5.8S rRNA (data not shown), indicating that it is defective in A_2_ and A_3_ cleavage, and truncated at the 5’-end.

To locate the 5’-end precisely we utilized reverse transcription (**Figure 2G**), which shows multiple 5’-ends between nucleotide 410 and 444. Mapping these positions onto the processosome structure demonstrates their location on the assembling body, in proximity to Rrp9 (orange spacefill in **Figure 2H**). Thus, removal of the first S1 domain of Rrp5 allows nucleases to access rRNA adjacent to Rrp9, supporting its binding near Rrp9 in early processosomes. Moreover, we also note the nearby crosslinking sites for Rrp5 (pink in **Figure 2H** (Lebaron *et al*., 2013)). Thus, deletion of Rrp5’s first S1 domain does not affect A_0_ cleavage but leads to defective A_2_ processing. Moreover, the A_0_-cleaved intermediate is subject to specific miscleavage or degradation at the 5’-end, perhaps because its destabilized. Finally, the data also confirm defects in 60S maturation steps, including A_3_ cleavage, as all 27S intermediates are strongly depleted, together with 25S and 5.8S rRNA. Moreover, polysome analysis demonstrates half-mers (data not shown), indicative of defects in 60S biogenesis.

Together, the protein-protein binding data, Rrp5 crosslinking and mapping of miscleavage sites all show that the first S1 domain in Rrp5 localizes Rrp5 to the Rrp36/Rrp9 complex on the assembling body, and that the disruption of this interaction leads to defects in assembly of the nascent body.

### Untethering Rrp5 from the body early in assembly leads to platform assembly defects

Above we have shown that early in 40S assembly Rrp5 binds to Rrp36/Rrp9, and that this interaction promotes productive assembly of the body. We hypothesized that this interaction also precludes the premature binding of Rrp5 to the platform, where it can promote misassembly and a dead-end assembly intermediate. Thus, we next asked whether untethering Rrp5 from the body leads to defects in platform assembly. To assess misassembly, we purified 40S ribosomal subunits from cells expressing either full-length Rrp5 or Rrp5_ΔS1 and then used semi-quantitative mass-spectrometry to assess the occupancy of ribosomal proteins (**Figure 3A, Supplemental Table S1**). Three biological replicates were analyzed for each sample and the spectral counts for each ribosomal protein were normalized by the total number of counts from small ribosomal subunit proteins. This analysis identified all proteins from the small subunit, although for Rps26 and Rps27, two of the smallest proteins, the summed average from both samples contained 35 and 31 peptides, respectively, limiting the significance analysis.

**Figure 3.**
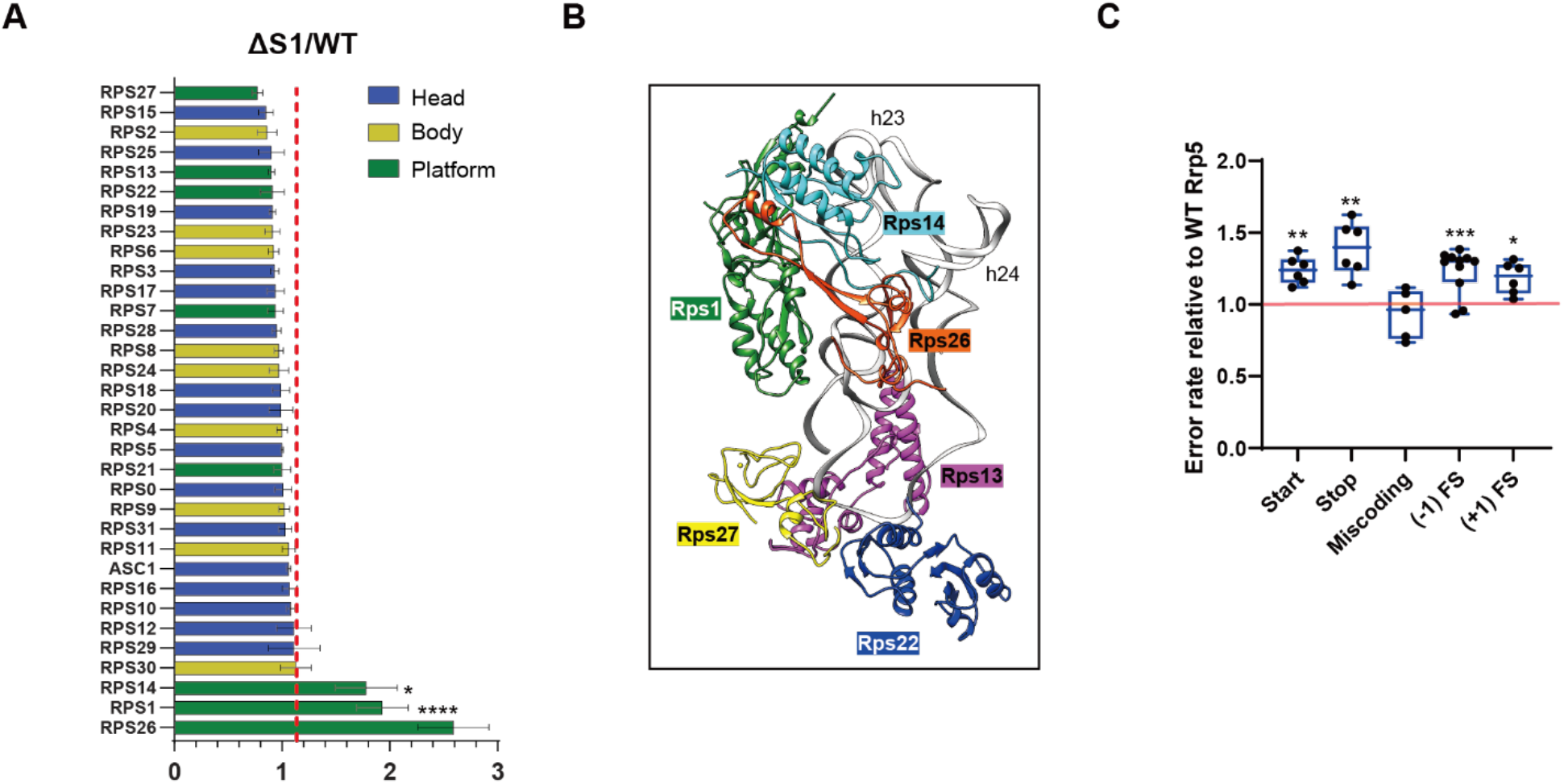
Disrupting the binding of Rrp5 and Rrp36 at the body leads to defective platform assembly and translation defects. A. Abundance of ribosomal proteins in 40S subunits purified from Rrp5_ΔS1 cells relative to WT cells determined by mass spectrometry. The red line indicates the average abundance of all ribosomal proteins. The data are averages from 3 biological replicates and error bars indicate the SEM. *: p <0.1 and ***: p<0.001 by unpaired t-test. B. Detail of the assembled platform in mature 40S ribosomes, showing the location of Rps1(Green), Rps13(Magenta), Rps14(Cyan), Rps22(Blue), Rps27(Yellow) and Rps26(Orange). From PDB ID 6ZCE. C. The effects from the Rrp5_ΔS1 mutation on start codon recognition, decoding, stop codon recognition, and programmed frame shifting (−1 and +1) was assayed using dual-luciferase reporters in a Gal::Rrp5 strain supplemented with plasmids encoding wild type or mutant Rrp5. Shown are the relative error rates of the Rrp5_ΔS1 samples relative to WT Rrp5. The data are the average of at least 5 biological replicates and error bars indicate the SEM. *: p <0.1, **: p<0.01 and ***: p<0.001 by unpaired t-test.

Comparing the occupancy of each ribosomal protein in the two samples, it becomes apparent that the majority of Rps are unchanged between the two samples. Intriguingly, Rps1, Rps14 and Rps26, which bind each other directly, are all significantly enriched in ribosomes from Rrp5_ΔS1 cells. This could arise from premature binding of these three proteins, which are depleted in 80S-like ribosomes (Ghalei et al., 2017), the last stable precursor to 40S subunits. In contrast, Rps27 is substantially depleted in the 40S subunits from Rrp5_ΔS1 cells. Moreover, its direct binding partners Rps13, and Rps22 are also depleted, albeit not as much as Rps27. Together, Rps27, Rps13 and Rps22 make up the base of the platform, while Rps1, Rps14 and Rps26 form the top of the platform. Thus, these data provide evidence for platform misassembly in ribosomes from Rrp5_ΔS1 cells, leading to depletion of components at the bottom (**Figure 3B**, Rps27, Rps13 and Rps22) and enrichment of components at the top (Rps1, Rps14 and Rps26).

Next, we asked if these defects in platform assembly led to defects in translation. The platform harbors the mRNA exit channel, which monitors the context of the start and stop codons and the E-site, from which tRNAs exit the ribosome. Weakened tRNA binding to this site increases frameshifting(Devaraj et al., 2009; Marquez et al., 2004), and the codon context is important for selection of both the start and stop codons. To test if ribosome populations have altered the fidelity of these steps, we have used previously described fidelity assays that depend on mistranslation in order to produce firefly luciferase(Ghalei et al., 2017). To control for differences in translational capacity, firefly luciferase activity is normalized against renilla luciferase, encoded on the same plasmid. Comparing the error rate in Rrp5_ΔS1 cells to the error rate in wt cells, we show that deletion of the first S1 domain in Rrp5, which allows for premature location of Rrp5 to the platform, leads to defects in start and stop codon selection as well as reading frame maintenance, as expected from defects in platform assembly. Decoding is not affected, showing that the A-site is properly formed (**Figure 3C**).

Thus, these data support a model where tethering Rrp5 to the body early in assembly, via the interaction between Rrp5 and Rrp36 serves to prevent misassembly resulting from premature binding of Rrp5 to the platform.

### Inactivation of the Has1 ATPase activity blocks the first rRNA processing step

The data above show that Rrp5 has two distinct locations on assembling 40S ribosomes: early in assembly it binds to Rrp9/Rrp36 on the nascent body, while later in assembly it binds to Utp22 and Krr1 on the developing platform, where it has also been visualized in available cryo-EM structures (**Figure 2H**,(Barandun *et al*., 2017; Du *et al*., 2020; Sun *et al*., 2017)). We next asked how Rrp5 is released from the nascent body to the platform. Previous work had shown that Rrp5 bound the DEAD-box ATPase Has1(Khoshnevis *et al*., 2016). Furthermore, crosslinking had suggested the existence of a Has1 binding site adjacent to an Rrp5 binding site on the assembling body (cyan in **Figure 2H**, (Bruning et al., 2018; Gnanasundram et al., 2019; Lebaron *et al*., 2013)). We therefore hypothesized that the RNA-dependent ATPase Has1 functions to relocate Rrp5 from the body to the platform, thereby integrating the assembly of these two subdomains.

Rrp5 and Has1 (and the ATPase Prp43) are the only assembly factors required for assembly of *both* ribosomal subunits. To dissect the role of Has1 in 40S assembly, we utilized two inactive Has1 mutants, Has1_T230A and Has1_H375A (**Figure S1B**). While Has1_T230A does not retain ATPase activity, Has1_H375A has futile ATPase activity that does not lead to effective remodeling (Rocak et al., 2005). As a result, Has1_T230A is a lethal mutation, while Has1_H375A shows severe growth defects (**Figure S1D, E)**, consistent with previous data (Dembowski et al., 2013; Rocak *et al*., 2005). Previous work had shown that these mutants could provide for Has1’s roles in 60S assembly, while 40S assembly required its ATPase activity (Dembowski *et al*., 2013; Gnanasundram *et al*., 2019).

First, we determined which of the early 40S rRNA processing steps was affected by Has1 inactivation. Previous work indicated that the A_2_ step was blocked in Has1 mutants (Dembowski *et al*., 2013), but did not specifically examine the preceding steps. We therefore used Northern blotting, combined with primer extension analysis to probe the effects from Has1 inactivation on rRNA processing (**Figure S3B**). The data demonstrate depletion of the A_2_ cleavage products, 20S rRNA and 27SA_2_ RNA. Moreover, lack of accumulation of 22S or 21S rRNA indicated that the preceding A_0_ and A_1_ cleavage steps were also disrupted (**Figure S1A**). To test this more directly, we employed primer extension analysis (**Figure S3C**), which showed strong inhibition of A_0_ cleavage, with a slightly stronger effect from the T230A mutant, consistent with the observed stronger growth phenotype from that mutant. Thus, the Has1 ATPase activity is required for the first rRNA processing step, A_0_ cleavage. This observation is consistent with our hypothesis that Has1 relocates Rrp5 away from the Rrp36/Rrp9 complex, as Rrp5 is already located at the platform in the structures of intermediates cleaved at the A_0_ site(Barandun *et al*., 2017; Du *et al*., 2020; Sun *et al*., 2017).

### Has1 relocates Rrp5 from the assembling body to the platform

Next, we asked whether deletion of the first S1 domain in Rrp5 rescued the growth and rRNA processing defects of the Has1_H375A mutation, as expected if Has1 releases Rrp5 from Rrp36. Thus, we combined the Has1_H375A and Rrp5_ΔS1 mutations and measured their effects on yeast growth quantitatively. These experiments demonstrate that deletion of the first S1 domain in Rrp5 (Rrp5_ΔS1) nearly completely rescues the growth defect from the Has1_H375A mutation (**Figure 4A**), providing strong genetic support for a role of Has1 in repositioning Rrp5 from the assembling body to the platform very early in 40S subunit assembly.

**Figure 4.**
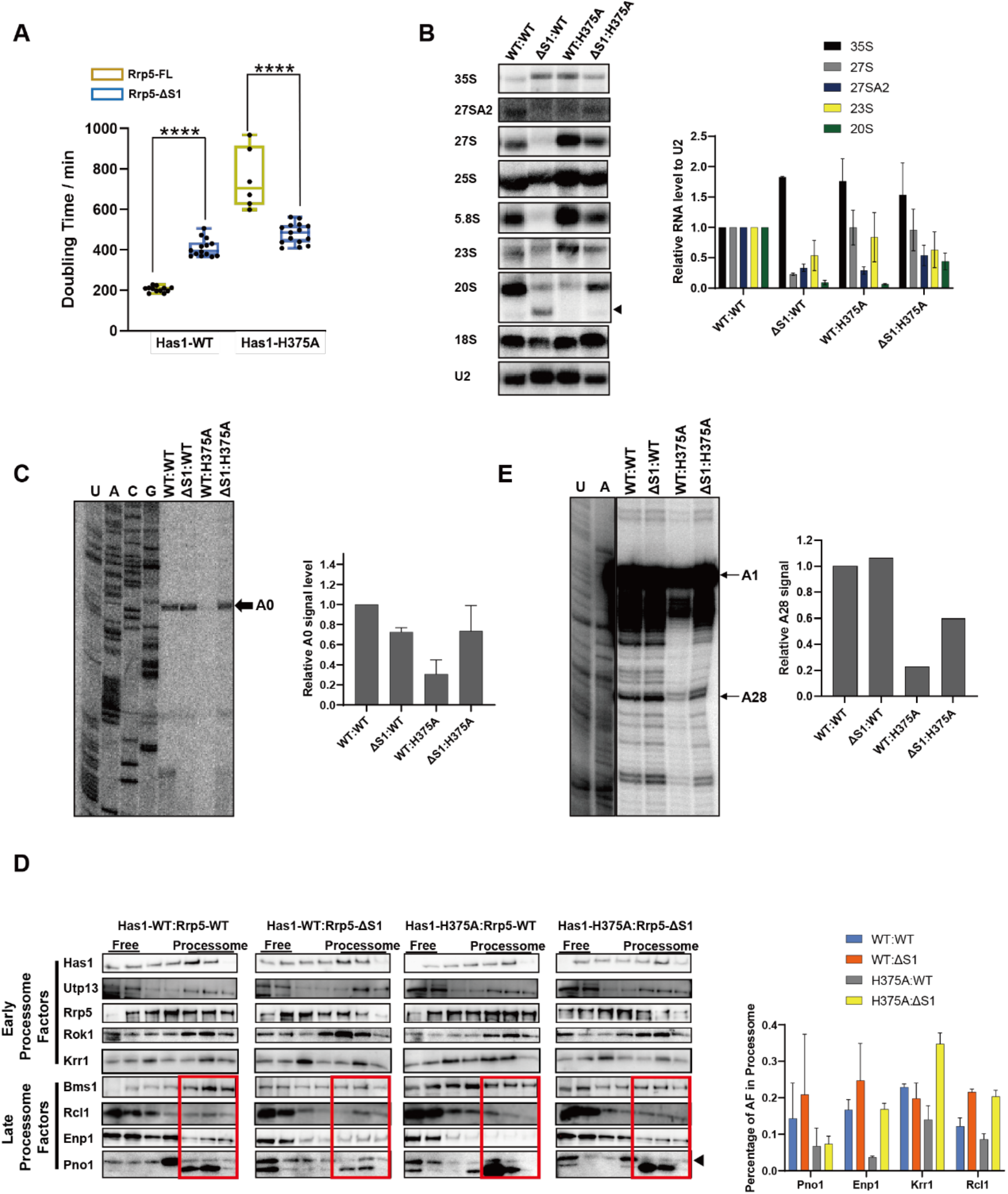
Rrp5-S1 deletion rescues Has1 inactivation. A. Doubling times of Gal:Has1;Gal:Rrp5 cells supplied with plasmids encoding Has1 WT or Has1 H375A and Rrp5 WT or Rrp5_ΔS1. Significance was tested using an unpaired t-test. ****, p<0.0001. n≥6 B. Northern blot analysis of rRNA processing intermediates from Gal:Has1;Gal:Rrp5 cells expressing plasmid-encoded Has1 and Rrp5 as in A. A degradation or mis-cleavage product in Rrp5_ΔS1 is indicated with arrow on the right. U2 serves as loading control. For the quantifications rRNA levels were normalized to U2 and to the sample with wild type Has1 and Rrp5. The data are averages from 3 biological replicates and error bars show standard deviation of the mean. C. Analysis of A_0_ cleavage by reverse transcription of total RNA extracted from Gal:Has1;Gal:Rrp5 depleted cells expressing plasmid-encoded protein as in A. Shown below is the quantification of A_0_ cleavage band, normalized to the full extension band (not shown). The data are averages from at from 2 biological and 2 technical replicates and error bars show standard deviation of the mean. D. Western blots of 10-50% sucrose gradient fractions from Gal:Has1;Gal:Rrp5 cells expressing plasmid-encoded Has1 and Rrp5 as in A. Free, 40S or 90S fractions are indicated with blue, red or green boxes, respectively. The band corresponding to Pno1 is indicated with an arrow on the right. Shown below are quantifications of the data from two or more biological replicate experiments and error bars show standard deviation of the mean. E. Modification of A28 requires Has1 activity and is bypassed with Rrp5_ΔS1. Reverse transcription is used to probe the snR74-dependent 2’-O methylation of A28.

To test whether disruption of the interaction between Rrp5 and Rrp36 also rescued the RNA processing defects from Has1 inactivation, we carried out Northern analysis. The defects in A_2_ cleavage in the Has1_H375A mutant (20S and 27SA_2_) depletion are rescued by the Rrp5_ΔS1 truncation (**Figure 4B**, right two lanes). Similarly, the accumulation of the aberrant 23S rRNA in the Has1_H375A background is reduced by the Rrp5_ΔS1 truncation, and primer extension demonstrates that A_0_ cleavage is restored (**Figure 4C**). Thus, disrupting the interaction between Rrp5 and Rrp36 at the nascent body rescues the growth and rRNA processing defects from the Has1_H375A mutation, providing strong evidence for a role of Has1 in separating Rrp5 and Rrp36.

### Inactivation of the Has1 ATPase leads to accumulation of Rrp36 in pre-40S ribosomes

Has1 is a member of the DEAD-box family of ATPases. These enzymes are RNA-dependent ATPases, which can disrupt very short RNA-RNA duplexes or RNA-protein interactions (Cordin *et al*., 2006; Jarmoskaite and Russell, 2014; Linder and Jankowsky, 2011). Thus, we assumed that to release Rrp5 from Rrp36, Has1 was acting on an RNA. The simplest model would be that Has1 releases Rrp36 from its RNA binding site, accounting for the lack of Rrp36 co-purification in all structurally characterized 40S assembly intermediates, as these are all later-forming intermediates.

To test if indeed Has1’s ATPase activity was used to release Rrp36, we carried out gradient sedimentation experiments in cells containing wild type Has1 or the inactive Has1_H375A. Notably, Rrp36 was accumulated in ribosome bound fractions (**Figure 5A)**. These data support a model where Has1 releases Rrp36 from very early pre-40S ribosomes, thereby enabling repositioning of Rrp5 from its interaction with the Rrp36/Rrp9 complex on the body to the platform. These data are further supported by mass-spectrometry analyses of assembly intermediates from wild type or Has1 mutant cells, which also show Rrp36 accumulation (see next section). Moreover, Rrp36 is a high-confidence interactor of Has1 in a genome-wide screen(Vincent et al., 2018).

**Figure 5.**
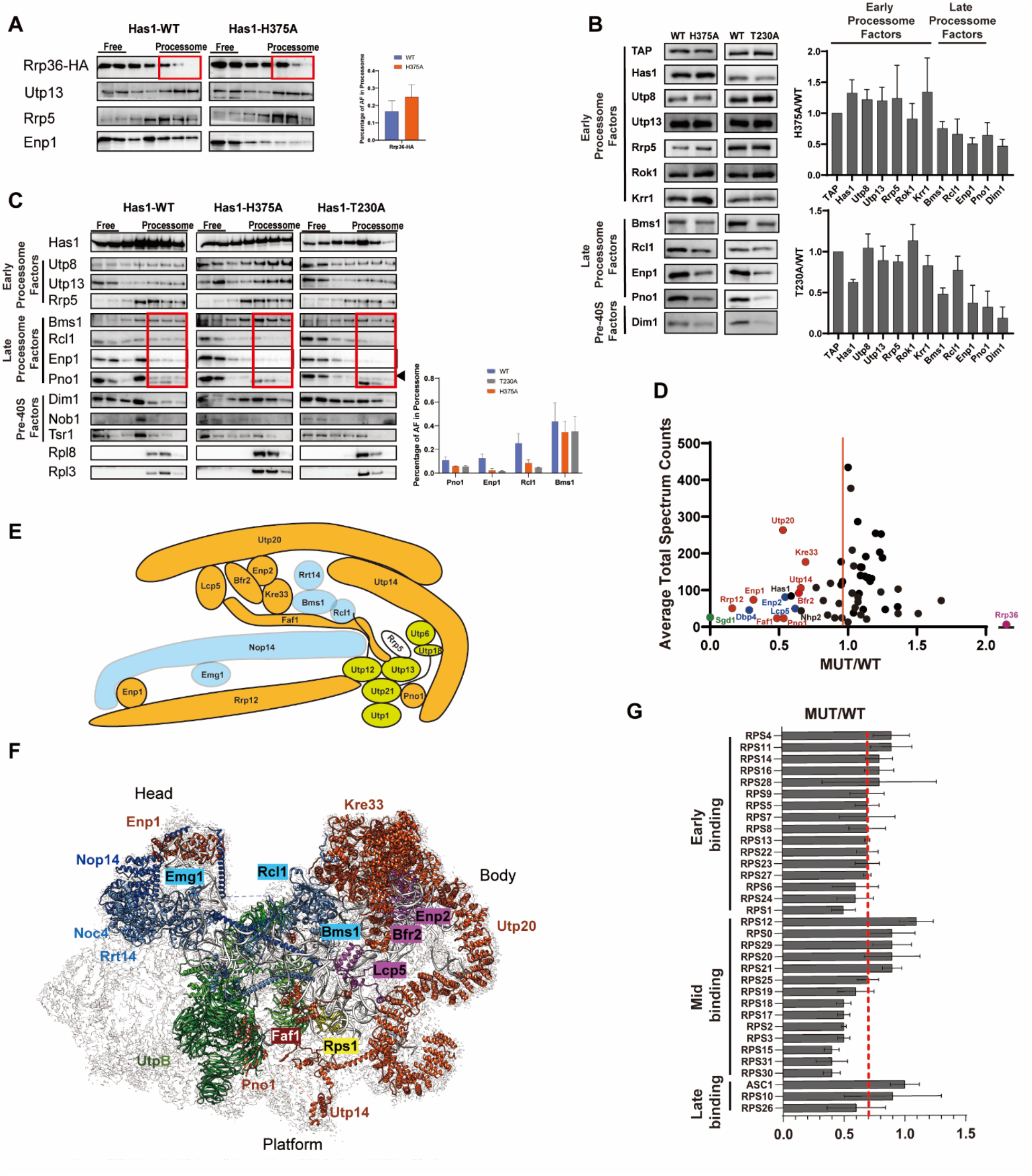
Assembly intermediates from Has1 mutant cells are defective in recruitment of late 90S pre-ribosome factors. A. Rrp36 accumulates in early processosomes after Has1 inactivation. Western blots of 10-50% sucrose gradients from Has1-depleted cells expressing plasmid-encoded Has1 WT or Has1_H375A. Free or processosome-bound Rrp36 was indicated B. Western blots of the Utp10-TAP elution from Has1 WT or mutant cell lysates. Utp10-TAP serves as loading control. Quantification of Western blots is shown on the right. Protein levels were normalized to Utp10-TAP levels and to the Has1 WT. The data are averages from at least 2 biological replicate experiments and error bars show standard deviation of the mean. C. Western blots of 10-50% sucrose gradients from Has1-depleted cells expressing plasmid-encoded Has1 WT or mutants. Free, 40S or 90S fractions were indicated with blue, red or green boxes, respectively. The band corresponding to Pno1 is indicated by the arrow on the right. Quantifications of the data from at least 2 replicate experiments indicate the distributions of late assembly factors in free, 40S or 90S fractions. The data are averages from at least 2 biological replicates and error bars show standard deviation of the mean. D. Abundance of assembly factors in Utp10-TAP elutions from Has1-H375A cells relative to wt cells determined by mass spectrometry. Late binding processosome factors are labeled in red, the Enp2/Lcp5/Bfr2/Dbp4 complex is shown in blue. Relative abundance (normalized to Utp10) was plotted against total spectral counts for each protein. E. Schematic representation of the network between late binding assembly factors (in orange or blue if or if not depleted in the mass spectrometry experiments in panel D) and UtpB components (green). Data are summarized from cryo-EM structures(Barandun *et al*., 2017; Cheng *et al*., 2017; Sun *et al*., 2017) and crosslinking mass-spectrometry (Chaker-Margot et al., 2017). F. Selected AFs mapped onto the pre-ribosome structure (PDB ID 6LQP). Depleted late-binding factors are shown in red, remaining late-binding factors are shown in blue, UtpB is shown in green. The depleted Rps1 is shown in yellow and the Enp2/Lcp5/Bfr2 complex is in purple. The locations of Dbp4 and Rrp14 are not known. G. Abundance of 40S ribosomal proteins in Utp10-TAP elutions from Has1-H375A cells relative to wt cells determined by mass spectrometry. The red line indicates the average abundance of all ribosomal proteins. The data in D and G are averages from 3 biological replicates.

### Inactivation of the Has1 ATPase activity blocks the recruitment of late processosome factors

To test whether Has1 ATPase-mediated release of Rrp36 and repositioning of Rrp5 to the platform led to larger changes in the assembling subunit, we purified assembly intermediates from yeast cells expressing wild type or inactive Has1 using a TAP-tag on the early processosome component Utp10, and then analyzed these intermediates using a combination of western analyses, mass spectrometry and DMS mapping. Utp10 is a UtpA component, and thus one of the earliest factors to bind nascent RNA(Perez-Fernandez et al., 2007), which dissociates soon after A_1_ cleavage(Cheng *et al*., 2020; Du *et al*., 2020). Thus, this strategy will purify early and late processosomes, but not intermediate or pre-40S subunits (**Figure 1C**).

Analysis of the TAP eluate using a collection of antibodies against processosome components demonstrates that assembly of early components such as the UtpA subunit Utp8, the UtpB constituent Utp13, Rrp5, or its binding partner Rok1, are not perturbed by Has1 inactivation (**Figure 5B**). In contrast, Pno1, Enp1 are not effectively recruited to pre-40S in either the Has1_T230A or the Has1_H375A mutant (**Figure 5B)**. These two AFs are part of a larger group of late binding AFs, whose binding requires transcription of the 3’-end of 18S rRNA (Chaker-Margot *et al*., 2015; Chen *et al*., 2020; Hunziker *et al*., 2019; Zhang *et al*., 2016), and a switch in the UtpB subcomplex (see below). Bms1 and Rcl1, which are also part of this group, are affected less consistently.

Because there was the possibility that the purification process might perturb intermediates, we sought to confirm this conclusion independently in an *in vivo* experiment, where we analyzed the sedimentation of these assembly factors with small subunit assembly intermediates. The late processosome intermediates sediment around 80S/90S. The sedimentation of the early processosomes is uncharacterized, but the late-binding factors provide about 1.2 MDa in mass, suggesting they should sediment between the 40S and 80S peaks. Later post-90S and cytoplasmic assembly intermediates sediment as 40S particles. Lack of ribosome recruitment manifests in sedimentation on top of the gradient. Consistent with the analysis of the purified intermediates, we observed increased sedimentation of Rcl1, Enp1 and Pno1 on top of the gradient, reflecting their reduced recruitment in the Has1 mutant strains. None of the other factors are significantly perturbed (**Figure 5C**). Thus, the Western analysis both of purified precursors and of fractionated lysates shows that Has1’s ATPase activity is required for the recruitment of at least some late assembly factors to the 40S processosome.

To test if recruitment of all late-binding processosome factors was equally affected by Has1 inactivation, or whether there was a subset of late-binding 90S AFs that was recruited independent of Has1 activity, we used mass-spectrometry to analyze assembly intermediates purified via Utp10 from cells containing wild type Has1 or Has1_H375A. Three biological replicates were analyzed for each sample, and peptide counts for each protein normalized to the bait, Utp10. This analysis demonstrates that the UtpA, UtpB, and UtpC subcomplexes, as well as the Mpp10 complex, and box C/D and H/ACA components are equally bound to the Utp10 assembly intermediates. The same is true for most other early-binding processosome factors, with the exception of the Bfr2/Enp2/Lcp5/Dbp4 subcomplex (**Figure 5D, Supplemental Table 2**). In contrast, many, but not all late-binding 90S factors are depleted, including Enp1, Utp14, Utp20, Kre33, Rrp12, as well as Pno1 and Faf1. These last two do not rise to statistical significance, because there are few peptides even in the wild type sample, indicating that neither protein performs well in the mass spectrometry, not unexpected given their small size and highly basic nature. In contrast, the late 90S factors Bms1, Rcl1, Emg1, Utp30, Nop14, Noc4 and Rrt14 are not affected. Notably, Rrp36 is the only factor that accumulates in pre-40S from Has1 inactive cells (**Figure 5D**), strongly supporting the notion that Rrp36 release requires Has1 activity (see above).

Recent biochemical and structural analyses have suggested that the recruitment of late-binding factors to the 40S processosome requires a conformational change within the UtpB subcomplex, which repositions the Utp12/Utp13 components relative to the others and the rest of the processosome (Hunziker *et al*., 2019). This forms the binding site for the late-binding assembly factor Pno1, explaining why Pno1 recruitment requires the UtpB switch ((Hunziker *et al*., 2019), **Figure 1D)**. Pno1 is depleted in the intermediates from the Has1_H375A cells. Moreover, all the other assembly factors that are depleted are also linked directly or indirectly to Utp12/Utp13 (**Figure 5E**): the structures show that Faf1 binds directly to Utp13, and Rrp12, which is not resolved in the available cryo-EM structures, crosslinks to Utp12. Thus, three of the 7 depleted late binding 90S factors (Pno1, Faf1 and Rrp12) directly contact Utp12 or Utp13, and are therefore expected to be sensitive to the UtpB rearrangement. The remaining ones are part of a network with these factors: Enp1 crosslinks to Rrp12; Utp14 binds directly to the UtpB component Utp6 as well as Pno1. Moreover, Utp14 crosslinks and binds to Utp20, which in turn connects to the Bfr2/Enp2/Lcp5 subcomplex and Kre33; these latter four proteins are further linked into the network via interactions with Faf1, which directly binds Utp13. Nonetheless, recruitment of Bfr2, Enp2 and Lcp5 does not depend on Kre33(Cheng *et al*., 2019), and these proteins are not assembling as part of the UtpB switch (Chaker-Margot *et al*., 2015; Chen *et al*., 2020; Hunziker *et al*., 2019; Zhang *et al*., 2016).

Together, the mass-spectrometry and Western analyses of the pulldowns from assembly intermediates, as well as the gradient analyses demonstrate that a subset of late-binding 90S assembly factors are depleted from assembly intermediates in Has1 inactive Has1_H375A cells. The depleted factors are all connected directly or indirectly to the Utp12/Utp13 components of the UtpB complex (**Figure 5E**), which change their position in the formation of late 90S pre-ribosomes, or Pno1, which binds their rearranged interface. Thus, the Has1 ATPase regulates assembly of the mature and active 90S processosome, likely by stabilizing the UtpB conformational switch.

### Deletion of the first S1 domain in Rrp5 rescues the processosome assembly defects arising from Has1 inactivation

Next, we tested whether the processosome assembly defects from the Has1_H375A mutation were rescued by deletion of the first S1 domain in Rrp5. For this experiment we carried out the gradient sedimentation experiment described above, monitoring the co-sedimentation of late processosome factors with pre-40S (both 90S and later 40S), or free factors. As before, early binding processosome factors are unaffected by either the Has1_H375A mutation, the Rrp5_ΔS1 truncation, or their combination (**Figure 4D**). Similarly, the Rrp5_ΔS1 truncation does not affect recruitment of any tested factors to the processosome. Moreover, as before (**Figure 4B-D)**, the Has1_H375A mutation impairs the recruitment of Pno1 and Enp1, so that they sediment as free factors. Importantly, combining the Rrp5_ΔS1 truncation with the Has1_H375A mutation rescues these defects, demonstrating that removal of the first S1 domain in Rrp5 rescues the late processosome assembly defects observed in the Has1 mutants.

These data further support a role for Has1 in the release of Rrp5 from Rrp36 in the body to the platform and suggest that the defects in assembly of the late-binding processosome factors and the resulting rRNA processing defects observed in the Has1 mutant arise from the lack of Rrp5 at the platform rather than Rrp36 remaining on the body. Nonetheless, it is also possible that without the stabilizing interactions from Rrp5, Rrp36 dissociates on its own.

### Rps1 incorporation is linked to the Has1-mediated repositioning of Rrp5

Analysis of the ribosomal protein content from the Utp10TAP pulldowns from wild type or Has1 inactive cells indicates that early-binding ribosomal proteins are overall depleted (mut/wt average = 0.7, **Figure 5G, Supplemental Table 3**). This is likely a reflection of the weaker binding of the proteins to earlier assembly intermediates as previously observed(Ferreira-Cerca et al., 2005), which results in them being disordered in the structures of the assembly intermediates lacking late factors (Chen *et al*., 2020; Hunziker *et al*., 2019). Notably Rps1 is the only early-binding ribosomal protein that is more depleted than the average ribosomal protein in the Has1 mutant cells (**Figure 5G**). This observation indicates that assembly of the body (where Has1 acts) is linked to assembly of the platform (where Rps1 binds). Notably, the observation that Rps1 binding is impaired when Rrp5 relocation is blocked parallels the observation that that Rps1 occupancy is increased when Rrp5 is constitutively relocated to the platform with the Rrp5_ΔS1 variant.

### Inactivation of the Has1 ATPase activity affects the binding site of the UtpB subcomplex and rRNA interactions of the late-binding assembly factors

The mass-spectrometry data demonstrate that the Has1 ATPase-mediated repositioning of Rrp5 leads to binding of a subset of the late-binding 90S processosome factors. Their binding requires a conformational switch in UtpB, which repositions Utp12 and Utp13 with respect to the rest of the UtpB complex and the assembling subunit (**Figure 1D**, (Hunziker *et al*., 2019)). Thus, we hypothesized that the Has1 ATPase-mediated release of Rrp36 and subsequent repositioning of Rrp5 to the platform enables the UtpB switch, thereby allowing for the assembly of the mature processosome. To further test this model and learn about conformational rearrangements in the assembling subunit, in particular those connected to the late-binding assembly factors and UtpB, we sought to use structure probing of the assembly intermediates purified from wild type or Has1_H375A cells (**Figure S4A**). Purified intermediates were exposed to DMS (or mock treatment), which methylates A and C residues that are not protected by base pairing or protein binding, before extracting and fragmenting the RNA for subsequent library generation and deep-sequencing(Huang and Karbstein, 2021; McGlincy and Ingolia, 2017). DMS-modified A and C residues are misread during reverse transcription, leading to mutations relative to the annotated rDNA sequence. DMS dependent changes in the mutational rate of A and C, but not U and G validate the procedure (**Table S4)**.

Sequencing reads in both the 5’-ETS and ITS1 regions, as well as in U3 demonstrate the successful purification and analysis of assembly intermediates that terminate at site A_3_ and are bound to the snoRNA U3 (**Figure S4B, C**). Moreover, the data also indicate that the samples are bound to U14 (snR128), whose abundance in the complexes approaches that of U3 (**Figure S4D**) and whose binding site is accessible in early pre-40S intermediates **(Figure S4E)**. The only other snoRNA that is enriched to some extent in the pulldown samples is snR74 (**Figure S4F**), which directs the methylation of A28. Notably, the abundance of snR74 was significantly reduced in the samples from Has1_H375A cells, suggesting that Has1 activity was required for binding of snR74 to pre-18S rRNA (**Figure S4F**). To test this hypothesis, we utilized reverse transcription to probe for the methylation at A28 imparted by snR74. As expected from the reduced occupancy of snR74 in the pulldowns from Has1_H375A cells, A28 methylation was reduced about 5-fold in Has1_H375A cells (**Figure 4E**), independently confirming the DMS MaPseq data.

A28 is located in close proximity to Rrp9, the Rrp5 crosslinking sites, and the miscleavage sites introduced by the Rrp5_ΔS1 deletion (**Figure 2H)**. We therefore hypothesized that the failure to recruit snR74 could arise from a steric block by Rrp5 when Has1 is inactivated. To test this hypothesis, we probed whether removal of the first S1 domain in Rrp5 rescues the A28 methylation defect observed in Has1_H375A cells. Indeed, Rrp5_ΔS1 rescues the A28 methylation defect that arises from Has1 inactivation (**Figure 4E)**, suggesting the recruitment of snR74 to nascent 40S subunits is also restored. These data support the model that snR74 recruitment requires the repositioning of Rrp5 from the body to the platform, possibly due to steric interference. Notably, A28 methylation is only partially rescued in the Rrp5_ΔS1; Has1_H375A cells, even though Rrp5_ΔS1 does not show a methylation defect. Finally, we note that these data strongly suggest a relatively late recruitment for snR74, after initial assembly of the subunit.

Together, the RNASeq analysis of these samples demonstrates that binding and release of U14 is independent of Has1 and occurs prior to the Has1-mediated release of Rrp36, while the recruitment of snR74, which methylates A28, is Has1-dependent, possibly because of Rrp5/Rrp36 blocking recruitment of the snR74 RNP and occurs only after initial assembly factors are already bound to the nascent subunit.

Analysis of 4 replicate experiments identified 80 high-confidence changes in the modification patterns of pre-rRNA and its bound U3 and U14 RNAs (**Figure 6A-D, Table S5)**. Changes in DMS accessibility are spread throughout the entire pre-rRNA. Notably, many of the residues that become protected in response to the Has1-mediated changes (shown in red or magenta in **Figure 6**) are directly interacting with ribosomal proteins (**Figure 6E**). This is consistent with the mass-spectrometry results that show reduced co-purification of ribosomal proteins (see above), the observation that ribosomal proteins bind more weakly early in assembly(Ferreira-Cerca *et al*., 2005), as well as the observation that ribosomal proteins are not resolved in assembly intermediates prior to the UtpB switch (Hunziker *et al*., 2019).

**Figure 6.**
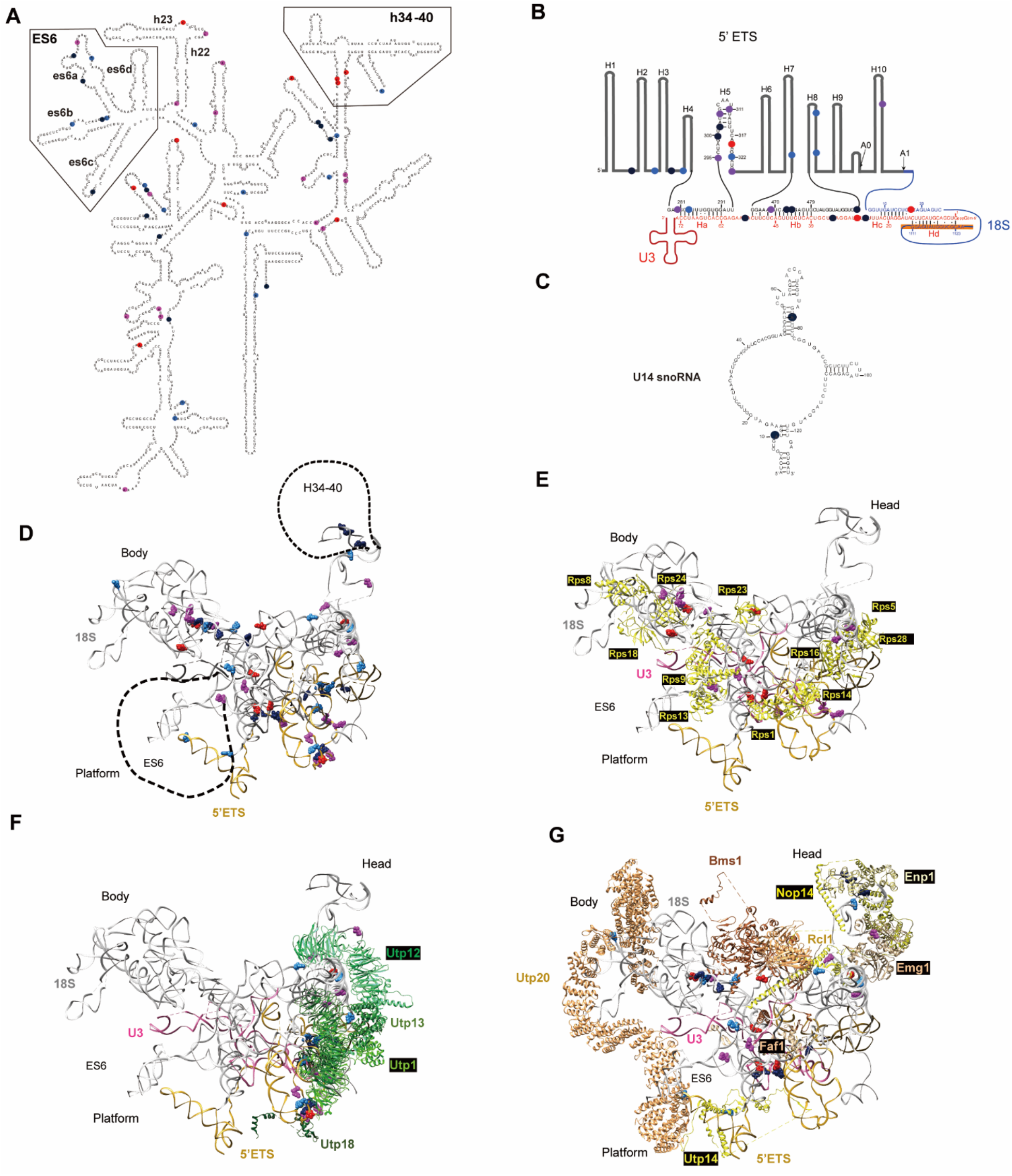
DMS-mapping shows that rRNA regions contacted by late processome factors are disturbed in the Has1 mutants. A-C. Nucleotides with differences in DMS accessibility between Has1 WT and the Has1_H375A mutant were mapped on the secondary structure of 18S rRNA (A), the U3/rRNA duplex (B) and U14 (C). Nucleotides that are more protected in the late processosomes from wt cells are shown in red or magenta (switched by at least two or just one bin, respectively). Nucleotides that become more exposed in the late processosomes from wt cells are shown in dark or light blue (switched by at least two or just one bin, respectively). Only statistically significant changes from 4 biological replicates (p_adj_<0.05, using a two-way ANOVA analysis) are shown. D. Changes in A and B were mapped onto the pre-ribosomal RNA (PDB ID 6LQP). 18S, and 5’ ETS are shown in gray and gold, respectively. E. Residues that are more exposed in mutant intermediates are located near ribosomal proteins. F. UtpB changes during the Has1-mediated change in assembly. Residues near UtpB that are becoming protected or exposed during the Has1 transition are shown. G. Changes in DMS accessibility near late binding assembly factors are shown. Late assembly factors depleted according to mass spectrometry are shown in orange, those that remain unchanged or accumulate in black.

Most of the remaining residues that become protected as a consequence of Has1 activity are contacted by Utp12 and Utp13, the two UtpB components that are repositioned in the UtpB switch (**Figure 6F**). Moreover, many of the residues that are *exposed* in the Has1-mediated conformational change (shown in dark or light blue in **Figure 6F**) are contacted by Utp12, the shifting UtpB components Utp1 and Utp21, as well as Utp18. Thus, residues directly contacted by UtpB make up a large fraction of the residues that change in the Has1-mediated transition, with some becoming protected and others exposed. Thus, these data provide direct support for our hypothesis that Has1-mediated release of Rrp36 and repositioning of Rrp5 enables the UtpB switch.

The final group of residues that change (becoming protected or exposed) as a result of Has1 activity are contacted by the late-binding assembly factors (**Figure 6G**). These changes affect both late-binding factors depleted in our mass-spectrometry (such as Enp1, Utp20, Utp14 and Faf1), as well as others that are not (Bms1, Rcl1 and Nop14), and suggest strongly that even the factors that are not depleted rearrange in the Has1-mediated change. Consistently, we note that by Western analysis of gradient centrifugation experiments Rcl1 and Bms1 are depleted from early pre-ribosomes and accumulated in the free fraction (**Figure 5B-C**) in the Has1_H375A and the Has1_T230A cells, perhaps reflecting loosed binding in those cells, that is below the threshold we apply in our mass spectrometry experiments.

In addition to the residues analyzed and discussed above, which are in regions of the nascent 40S that are visualized in the processosome structures, two regions with a significant number of differentially accessible residues are not resolved in the cryo-EM structures of the processosome: the ES6 (expansion segment 6) region, a eukaryote-specific structure, which pins the platform to the body, as well as h34-h40, which comprise much of the nascent subunit’s head (**Figure 6A)**. Notably, ES6, in particular ES6A and ES6D, are more protected in the molecules from Has1 mutants, consistent with premature folding or misfolding of these areas. Lcp5, which is depleted in intermediates from the Has1 mutant cells, blocks the formation of ES6D(Cheng *et al*., 2020; Du *et al*., 2020). We thus suggest that the premature formation of ES6D is due to the loss of Lcp5. In contrast, residues in h34-40 are more exposed in the samples from Has1 mutant cells, consistent with their folding after Has1-mediated progression in the assembly cascade.

In summary, the DMS-MaPSeq data provide strong independent confirmation for changes in the UtpB subcomplex arising from Has1 activity, and in particular its Utp12 and Utp13 components. These changes are propagated to other late-binding assembly factors as well as the early-binding ribosomal proteins, stabilizing the incorporation of the ribosomal proteins after the Has1-mediated release of Rrp36 and repositioning of Rrp5, and mediating the recruitment of and repositioning of the late-binding assembly factors.

### Genetic interactions with Has1_H375A support its role in the UtpB switch

As described above, the biochemical and structural analyses of assembly intermediates, both purified and within cells indicate that the Has1 ATPase-mediated release of Rrp36 from early pre-40S assembly intermediates and the resulting repositioning of Rrp5 from the body to the platform also promotes the UtpB switch. To further test this model, we carried out two sets of genetic experiments.

If Has1 promotes the UtpB switch and the assembly of the late processosome, then mutations that destabilize the processosome are expected to be synthetically sick with the partially active Has1_H375A. To test this prediction, we used the 90S structures to produce mutations in late-binding processosome factors (Faf1, Emg1), which are predicted to weaken their binding by disrupting their interactions with the RNA (**Figure 7A-B**, left). These mutants were then tested in galactose-inducible/glucose repressible strains where both the endogenous copy of the test protein and Has1 can be depleted by growth in glucose. These strains are then supplemented with plasmids encoding wild type or mutant Has1 and either wild type or mutant Faf1 or Emg1, and doubling times measured quantitatively in continuous growth measurements. As expected if Has1 promotes the formation of the late processosome, the Faf1_R239A/R240A mutation and the Emg1_T127/R129/R132/T133A mutation are synthetically sick with Has1_H375A (**Figure 7A-B**, right).

**Figure 7.**
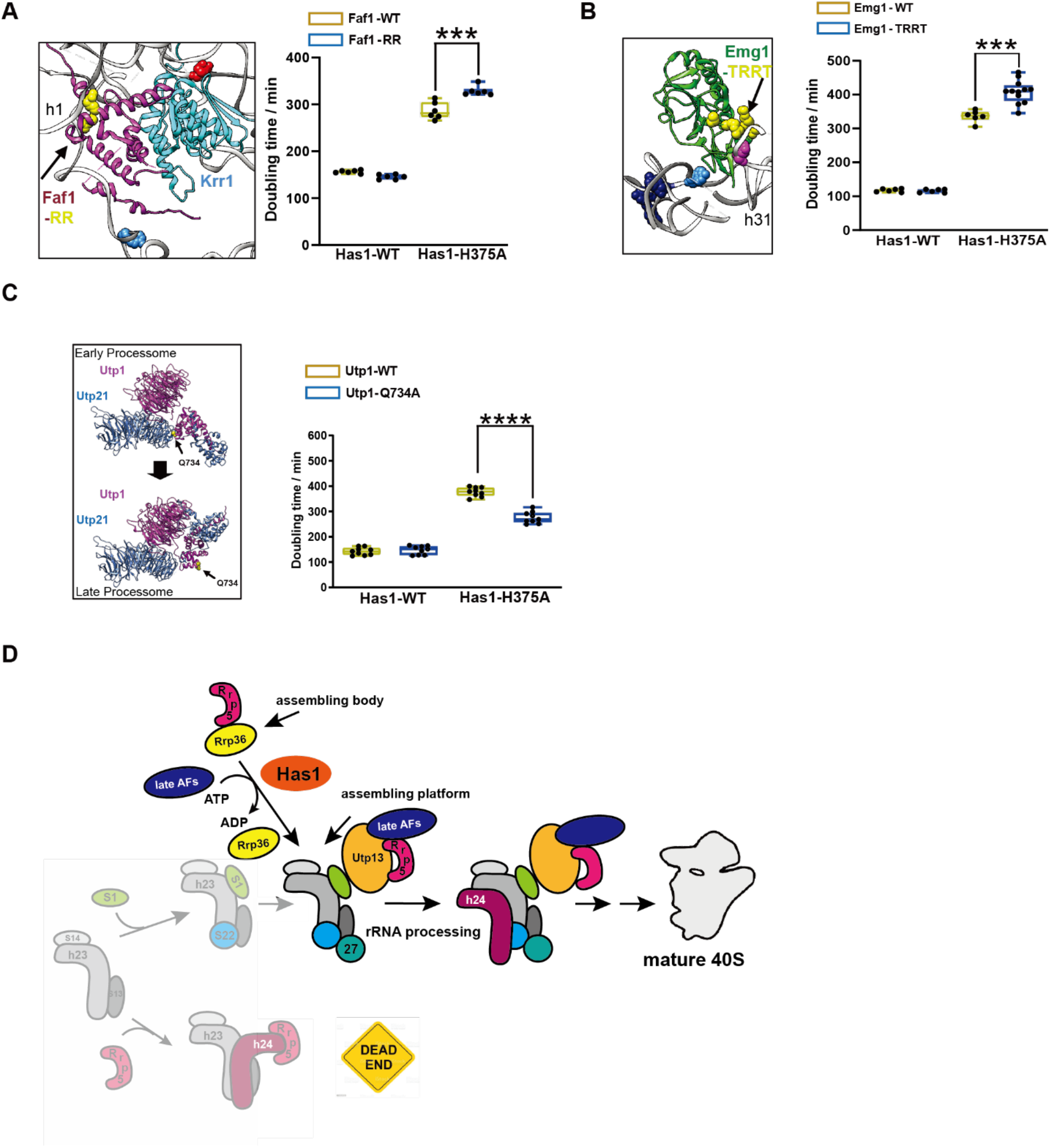
Promoting the UtpB conformational change genetically rescues Has1 inactivation. A. Left, the location of Faf1_RR (R239A/R240A) is indicated (from PDB ID 6LQP). Right, doubling times of Gal:Has1,Gal:Faf1 cells supplied with plasmids encoding Has1 WT or Has1_H375A or Faf1 WT and Faf1_RR. Significance was tested using an unpaired t-test. ***, p<0.001. n≥7 B. Left, the location of Emg1_TRRT (T127A/R129A/R132A/T133A) is indicated (from PDB ID 6LQP). Right, doubling times of Gal:Has1,Gal:Emg1 cells supplied with plasmids encoding Has1 WT or Has1_H375A or Emg1 WT and Emg1_TRRT. Significance was tested using an unpaired t-test. ***, p<0.001. n≥6 C. Left, the UTP-B confirmational switch repositions Utp1_Q734 from an interaction with Utp21 in 90S structures (PDB ID 6ND4) to the solvent in late 90S structure (PDB ID 6LQP). Utp1_Q734 is indicated in yellow space fill. Right, doubling times of Gal:Has1,Gal:Utp1 cells supplied with plasmids encoding Has1 WT or Has1 H375A and Utp1 WT, or Utp1 Q734A. Significance was tested using an unpaired t-test. ****, p<0.0001. n≥10 D. Model for the roles of Rrp36 and Has1 in preventing premature Rrp5 binding to the platform to ensure its proper assembly. By tethering Rrp5 to the assembling body (top) early in assembly, its premature binding to the platform (bottom, shaded) is prevented, thus avoiding the folding funnel that leads to the dead-end. By releasing Rrp36 from the assembling body, Has1 allows Rrp5 to relocate to the platform (middle), where it stabilizes Utp13 in the switched structure, thereby allowing for the UtpB switch and the recruitment of late assembly factors (late AFs, blue). This enables A_0_ processing, thereby ensuring that only correctly assembled processosomes are activated for A_0_ cleavage.

Next, we tested the opposite prediction. If Has1 promotes the formation of the late processosome via the UtpB switch, then mutations that destabilize the early structure should partially rescue the Has1_H375A mutation. For this experiment, we evaluated the interfaces between the individual UtpB components in the early and late processosome and designed the Utp1_Q734A mutation. Q734 interacts with Utp21 only in early, but not late processosomes (**Figure 7C**, left). Indeed, Utp1_Q734A partially rescues the Has1_H375A mutation (**Figure 7C**, right), providing strong genetic support for the model that the Has1 ATPase activity supports the conformational switch in UtpB, which enables the recruitment of late processosome factors, and ultimately rRNA processing at site A_0_.

## Discussion

### Has1 releases Rrp36 allowing Rrp5 to reposition from the assembling body to the platform

Our biochemical analysis demonstrates that inactivation of the DEAD-box ATPase Has1 impairs the assembly of late-binding components of the 90S processosome, and thereby blocks the first rRNA processing step, 100 nt upstream of the mature 5’-end of 18S rRNA. Deletion of the first S1 domain in Rrp5 (Rrp5_ΔS1), an early-binding processosome component that interacts with Has1 (Khoshnevis *et al*., 2019; McCann et al., 2015; Tarassov et al., 2008), nearly completely rescues the growth, processosome assembly and rRNA processing defects observed in cells containing inactive Has1. Biochemical analyses with purified recombinant components demonstrate that the interaction partner for this subdomain of Rrp5 is the early assembly factor Rrp36. Thus, these data suggest that Has1 functions by separating Rrp5 and Rrp36. Gradient centrifugation coupled with Western analysis, as well as mass-spectrometry of purified intermediates both show an accumulation of Rrp36 in early pre-40S intermediates upon Has1 inactivation, suggesting very strongly that Has1 releases Rrp36 from early processosomes (**Figure 7D**). This model is consistent with the absence of Rrp36 in all structures or pulldowns of previously characterized assembly intermediates, as these are subsequent to the novel intermediate implicated here. Yeast two-hybrid analyses, supported by pulldowns, have shown that Rrp36 and Rrp5 are part of a larger substructure on the body of the assembling subunit, which also includes the U3 snRNP component Rrp9 (Clerget *et al*., 2020). Moreover, previous crosslinking data(Lebaron *et al*., 2013), rRNA miscleavage near the Rrp9 binding sites in the Rrp5_ΔS1 deletion, and the inability to recruit snR74 to this region in the Has1 mutant all support binding of Rrp5 to the assembling body early in 40S maturation. In contrast, biochemical experiments herein and previous cryo-EM structures (Cheng *et al*., 2020; Du *et al*., 2020; Sun *et al*., 2017) demonstrate that in later assembly intermediates Rrp5 binds on the assembling platform, where it interacts with Utp13, Utp22 and Krr1. Thus, these data strongly suggest that Rrp5 repositions during 40S maturation from a location on the assembling body, where it is bound to Rrp9/Rrp36, to the assembling platform, where is binds Utp13, Utp22 and Krr1. More importantly, the data demonstrate that Has1 regulates this repositioning by releasing Rrp36 from early assembly intermediates, thereby freeing Rrp5 up to relocate (**Figure 7D)**.

### Assembly factors can bias the folding landscape to avoid misfolding funnels

A large body of previous work has demonstrated that ribosome assembly proceeds via parallel pathways(Cheng *et al*., 2019; Cheng *et al*., 2020; Davis *et al*., 2016; Du *et al*., 2020; Mulder *et al*., 2010; Sanghai *et al*., 2018; Sashital *et al*., 2014). While these can speed up folding and render it more robust (Bedard et al., 2008; Fersht et al., 1994), *in vitro* analyses have demonstrated that some of pathways are dead-ends(Mulder *et al*., 2010). How these are avoided *in vivo* remained unclear until now.

While dead-end intermediates are typically not well-studied in vivo because they tend to be degraded, previous structural analyses of 40S assembly intermediates provide a unique opportunity to study one such intermediate, stable in starved cells and promoted by the premature location of Rrp5 at the platform: Comparison of numerous structures of assembly intermediates shows that all are nearly identical, except at the platform, where intermediates from wild type yeast show an assembling platform bound to Rps13/uS15, Rps14/uS11, Rps1 and Rps22/uS8. Notably, h24 is not yet formed. Rrp5 is located near the platform, held by protein-protein interactions with Utp13, Utp22 and Krr1. No interactions with rRNA are observed. In contrast, in structures from starved yeast, Rrp5 prematurely stabilizes and misorients h24, blocking the assembly of Rps22. These intermediates cannot convert to later assembly intermediates and thus represent a dead-end(Barandun *et al*., 2017; Du *et al*., 2020). Thus, premature binding of Rrp5 to the h24 on the platform leads to its misassembly and blocks further maturation (**Figure 1A&B**). How unstarved cells largely avoid this misassembly pathway remained unclear from these structures.

Herein, we show that premature binding of Rrp5 to the platform is blocked by its initial localization to the body. This avoids the branch in the assembly pathway that leads to the dead-end intermediate (shaded part of **Figure 7D**), thereby avoiding the dead-end. Thus, Rrp36 blocks a folding funnel that can lead to a dead-end, or misfolding, simply by avoiding the entrance into that pathway, opening a different route instead, which is not encumbered by misassembly. This is a novel function for an assembly factor.

In addition, the initial localization of Rrp5 to the assembling platform sets up a point of regulation for the DEAD-box ATPase Has1, which enables the switch in Rrp5 localization. Disrupting the interaction between Rrp5 and Rrp36 with the Rrp5_ΔS1 truncation not just circumvents the Has1-dependent checkpoint, but also allows for premature binding of Rrp5 to the platform. Accordingly, in these yeast strains the platform is misassembled, leading to reduced occupancy of Rps27 and increased binding of Rps1, Rps14 and Rps26. The increased occupancy of Rps1, Rps14 and Rps26, which all bind each other might reflect premature binding of these proteins to the latest assembly intermediates. The resulting misassembled platform not only is structurally deficient, but also functionally impaired. The platform contains both the tRNA E-site, which is important for reading frame maintenance during translocation(Devaraj *et al*., 2009; Marquez *et al*., 2004), and the mRNA exit channel, which monitors the context of the start and stop codons. Correspondingly, ribosomes from Rrp5_ΔS1 yeast are defective in the identification of start and stop codons, as well as reading frame maintenance. They do not have defects in decoding. Thus, tethering Rrp5 away from the platform early in assembly via its interaction with Rrp36, biases the folding landscape towards pathways that enable proper platform assembly and avoid dead-end pathways.

### Has1-mediated Rrp5 repositioning enables the UtpB conformational switch

In addition to its role in releasing Rrp5 from Rrp36, our data also demonstrate a role for the Has1 ATPase activity in the UtpB switch, which has been previously implicated in assembly of the processosome via recruitment of a group of late binding assembly factors (Hunziker *et al*., 2019). Biochemical, structural and genetic analyses demonstrate that a subset of these late-binding factors fail to assemble in the presence of inactive Has1, while the remaining ones fail to reposition. Moreover, destabilizing the late processosome produces synthetic genetic interactions, while destabilizing the early processosome partially rescues the inactivation of Has1. Thus, these data show that Has1 activity can be partially substituted by promoting the UtpB switch. How is Rrp5 repositioning and its role in proper assembly of the platform linked to the UtpB switch?

The UtpB switch reorients Utp12 and Utp13 with respect to the other UtpB components and the rest of the processosome(Hunziker *et al*., 2019). This enables the binding of the top of the decoding site helix, h44, to these two components, thereby explaining why transcription of all of 18S rRNA is required for the UtpB switch (Chaker-Margot *et al*., 2015; Chen *et al*., 2020; Hunziker *et al*., 2019; Zhang *et al*., 2016). Notably, the foot of h44 binds Utp22, which also interacts with Rrp5 and Utp13. This interaction with Rrp5 only occurs in the correctly assembled intermediates, as a different Utp22•Rrp5 interface is observed in misassembled structures. Finally, one of Rrp5’s S1 domains is adjacent to Utp13 in the switched structure. Thus, relocation of Rrp5 to the platform will stabilize UtpB in its switched structure, by helping to position Utp13 directly and by linking Utp22 to Utp13, thereby prepositioning it for interaction with h44.

### Using energy to impose order on assembly

Taken together the data herein show that the interaction between Rrp36 and Rrp5 on the assembling body, establishes a point of regulation in 40S maturation, which prevents the premature binding of Rrp5 to the platform, thereby avoiding a misfolded platform during maturation. Because Rrp5 repositioning stabilizes a conformational switch in UtpB, which is required for assembly of the processosome, and the first rRNA cleavage step, this checkpoint ensures not just that rRNA misfolding during platform assembly is avoided, but also guarantees that misassembled intermediates cannot fully assemble processosomes and are therefore incapable of carrying out the necessary RNA processing steps. Finally, our data demonstrate that this checkpoint and thereby the assembly of the processosome is regulated in an ATPase-dependent manner by the DEAD-box protein Has1. Thus, Rrp36, Rrp5, the UtpB complex and Has1 cooperate to ensure the correct assembly of the 40S platform in an ATP-dependent manner.

The use of energy to order a process that might otherwise be random is reminiscent of the role of the Rab proteins in the secretory pathway. These small GTPases regulate membrane trafficking by establishing microdomains in the membranes along the secretory pathway, which serve as molecular markers and move cargo in a directional and targeted manner(Pfeffer, 2017). Order in the pathway is imposed via the localization of GTP exchange factors (GEF) and GTPase activating factors (GAP) by the Rabs themselves. Thus, GTP is utilized in this system to establish the membrane compartments that are then recognized by cargo as it moves along the secretory pathway, similar to the use of ATP by the DEAD-box protein Has1 in ordering assembly of the body and platform. More generally, DEAD-box proteins should be considered ATP-dependent molecular switches, similar to GTPases, rather than performing mechanical work such as progressive duplex unwinding(Jankowsky, 2011; Jarmoskaite and Russell, 2011; Khoshnevis *et al*., 2016).

### Linking the assembly of subdomains and subunits via Rrp5 and Has1

Our data, together with previous work(Gerus et al., 2010), indicate that Rrp5 initially localizes to the assembling body, before being relocated to the assembling platform in a Has1-dependent manner. Thus, Rrp5 could link the assembly of the platform to that of the body. Indeed, the compositional analysis of assembly intermediates stalled in the absence of Has1 demonstrates that Rps1, a platform component, is depleted. Moreover, we also show that when body and platform assembly are unlinked, via the Rrp5_ΔS1 truncation, the platform is misassembled. Thus, the data herein provide evidence how Rrp5, Rrp36 and Has1 link assembly of the body and platform.

In addition to linking assembly of the body and platform, Rrp5 and Has1 also link assembly of the small and large ribosomal subunits, as they are two out of three proteins required for assembly of both ribosomal subunits, and not just one or the other subunit. Together, these observations underscore not just the importance placed on balanced subunit assembly as previously indicated(Khoshnevis *et al*., 2019), but also suggest the importance of avoiding misassembly of individual subdomains.

## Materials and Methods

### Yeast strains and plasmids

*Saccharomyces cerevisiae* strains used in this study are listed in **Table S6**. Yeast strains were generated using standard recombination techniques and verified by Western blotting and/or colony PCR. Plasmids used in this study are listed in **Table S7**.

### Protein expression and purification

All proteins were expressed in *E. coli* Rosetta2 (DE3) cells (Novagen). Cells were grown at 37°C to OD_600_ of 0.6 in LB media supplemented with the appropriate antibiotic and then transferred to 18°C. Protein expression was induced by addition of 0.3 mM or 1 mM IPTG for pGEX-6-P3 or pET23/pSV272 harboring cells, respectively, and cultures were harvested after 18 hours.

Rrp5 FL and Rrp5_S1, Rrp5_S1/S2, Rrp5_S1-3, Rrp5_ΔS1, Rrp5_ΔS1-S2, Rrp5_S7-C and Rrp5_S8-C were purified as previously described(Khoshnevis *et al*., 2016; Khoshnevis *et al*., 2019; Young and Karbstein, 2011).

MBP-Rrp36 was purified using amylose resin (New England BioLabs) in MBP-binding buffer [200 mM NaCl, 50 mM HEPES/NaOH (pH 7.5), 5 % glycerol]. Protein was eluted in MBP-binding buffer supplemented with 20 mM maltose. The complex was further purified using a MonoQ ion exchange column (GE Healthcare) equilibrated with MBP-binding buffer. The protein was eluted with a salt gradient [1M NaCl, 30 mM HEPES/NaOH, 10 % glycerol and 2 mM ΔME over 20 column volumes. The protein was further polished using a Superdex S-200 gel filtration column (GE Healthcare) equilibrated in 200 mM NaCl, 20 mM HEPES/NaOH, 10 % glycerol and 1 mM DTT.

MBP-Krr1 was purified similarly to MBP-Rrp36 with minor modifications: The protein was eluted with a salt gradient [1M NaCl, 50 mM Tris-HCl pH 7.5, 5 % glycerol and 2 mM ΔME over 20 column volumes. The protein was further polished using a Superdex S-200 gel filtration column (GE Healthcare) equilibrated in 200 mM NaCl, Tris-HCl pH 7.5, 5 % glycerol and 1 mM DTT.

### In vitro protein interaction studies

3 μM of FL Rrp5 or Rrp5 fragments were mixed with 3 μM MBP-Rrp36 in 250 mM NaCl and 20 mM HEPES/NaOH (pH 7.5), 5 % glycerol, and preincubated on ice for 30 min before addition of 30 μl of equilibrated amylose resin. The mixture was incubated for 30 min at 4°C, flow-through was collected, resin washed and eluted with binding buffer supplemented with 20 mM maltose. For Rrp5-MBP-Krr1 interaction studies 3 μM of FL Rrp5 were mixed with 3 μM MBP-Krr1.

### Northern and reverse transcription analysis of rRNA processing and modification

Cells were grown in the presence of glucose to OD ∼ 0.6 and total RNA was isolated using the hot phenol method. rRNA processing intermediates were analyzed either by reverse transcription using primers or by Northern blotting using the following probes: 35S, 23S and 27SA_2_: probe 003 (between A_2_ and A_3_), 27S and 5.8S: probe 004 (3’ of 5.8S), 20S: probe 002 (between D and A_2_), 18S and 25S: probes 001 and 005, respectively (within the mature rRNA of interest) in **Table S8**.

### Sucrose density gradient analysis

Sucrose gradient fractionation of whole cell lysates and subsequent western blot analyses were carried out as described before (Strunk et al., 2012). To calculate the relative amount of AFs in each fraction peak, we quantified the signal in the free fraction (fractions 1 and 2), 40S fraction (fractions 4) and the 90S fraction (fractions 6 and 7), and then divided the individual peak signal by the sum of the free, 40S and 90S signals.

### Growth curve measurements

Cells were grown in YPD overnight, and then diluted into fresh YPD for 3-4 hours before inoculating into 96-well plates (Thermo Scientific) at a starting OD600 between 0.04 to 0.1. A Synergy.2 plate reader (BioTek) was used to record OD600 for 24 hours, while shaking at 30 °C. Doubling times were calculated using data points within the mid-log phase. Data were averaged from at least 6 biological replicates of 3 different colonies and 2 independent measurements. Statistical analyses for each measurement are detailed in the respective figure legend.

### TAP purifications

Ribosome assembly intermediates were purified using Utp10-TAP via IgG and calmodulin beads essentially as previously described(Strunk et al., 2011), except that for Western and mass-spectrometry analyses only the IgG step was carried out. Both steps were used for DMA MaPseq analysis.

### DMS MaPseq sample preparation, library preparation, data processing and analysis

The purified pre-ribosomes were treated at 30°C for 5 min with 1% DMS (Sigma-Aldrich) in the presence of 80mM HEPES pH7.4, 50mM NaCl, 5mM Mg(OAc)_2_, 0.2uM RNaseP RNA.

DMS reactions were stopped by addition of 0.4 vol quench (1M β-ME, 1.5M NaOAc, pH 5.2) and purified using phenol chloroform precipitation. DMS MaPseq RNA libraries were prepared as previously described (ref, haina’s new paper), and analyzed by paired-end deep sequencing on the Illumina NextSeq 500 platform. Processing and analysis of the DMS MaPseq data followed the pipeline from GitHub code (https://github.com/borisz264/mod_seq) and as previously described (Huang and Karbstein, 2021). At least three biological replicates were collected for each sample. Control experiments demonstrate that DMS increases the mutational rate of A and C, but not G and U, as expected because DMS does not modify U, and because modification of G does not introduce a mutation upon reverse transcription. Raw mutational rates are summarized in **Table S1**. All significantly changed residues are listed in **Table S5**. To visualize changes in DMS accessibility we sorted the data into 5 bins, based on the normalized difference in the mutational rate with and without DMS: 0-0.5, 0.5-1, 1-2, 2-4, above 4; Next, we compared the bin assignment for each nucleotide in cells with wild type or mutant Has1. Residues that are changing one bin are shown in magenta (protected in wt) or light blue (exposed in wt), while those changing by two or more bins are shown in red or dark blue, respectively.

### Ribosome purifications

Ribosomes were purified as described(Collins *et al*., 2018). In brief, seeded 1-liter cultures were harvested at OD600 1.5–1.7 and flash-frozen in ribosome buffer (20 mM Hepes/KOH, pH 7.4, 100 mM KOAc, and 2.5 mM Mg(OAc)2) supplemented with 1 mg/ml heparin, 1 mM benzamidine, 2 mM DTT, and protease inhibitor cocktail (Sigma-Aldrich). Frozen cells were lysed by grinding to powder in a mortar and pestle. Yeast lysates were clarified and layered over a 500 µl sucrose cushion and centrifuged in a Beckman TLA 110.1 at 70,000 rpm for 70 min. Pelleted ribosomes were resuspended in high-salt buffer (ribosome buffer, 500 mM KCl, 1 mg/ml heparin, and 2 mM DTT) and again layered over a 500 µl sucrose cushion and centrifuged at 100,000 rpm for 100 min. The ribosome pellet was resuspended in subunit separation buffer (50 mM Hepes/KOH, pH 7.4, 500 mM KCl, 2 mM MgCl2, 2 mM DTT and 1 mM puromycin (Sigma-Aldrich). Subunits were isolated by loading onto 5–20% sucrose gradients (50 mM Hepes/KOH, pH 7.4, 500 mM KCl, 5 mM MgCl2, 2 mM DTT and 0.1 mM EDTA) and centrifuged at 30,000 rpm for 8 h. Finally, 40S subunits were concentrated, buffer exchanged into ribosome storage buffer (ribosome buffer with 250 mM sucrose and 2 mM DTT), flash frozen and stored at −80°C.

### Translational fidelity by luciferase assays

Translational fidelity was measured using the Dual-Luciferase® Reporter Assay (Promega) as previously described (Ghalei et al., 2017). Briefly, cells were grown to mid-log phase, 2ml of cells were pelleted, washed and flash frozen. Cells were resuspended in 1ml Passive lysis buffer. Firefly and renilla signals were measured sequentially by addition of 20µl Luciferase Assay Reagent II and 20µl Stop&Glo Reagent with 10µl lysate. For each sample, Firefly activity was first normalized to renilla activity, before normalizing the firefly/renilla ratio for each mutant to that for wild type. Data were derived from at least 6 biological replicates, with 3 technical replicates each.

## Supplemental Material to

**Figure S1.**
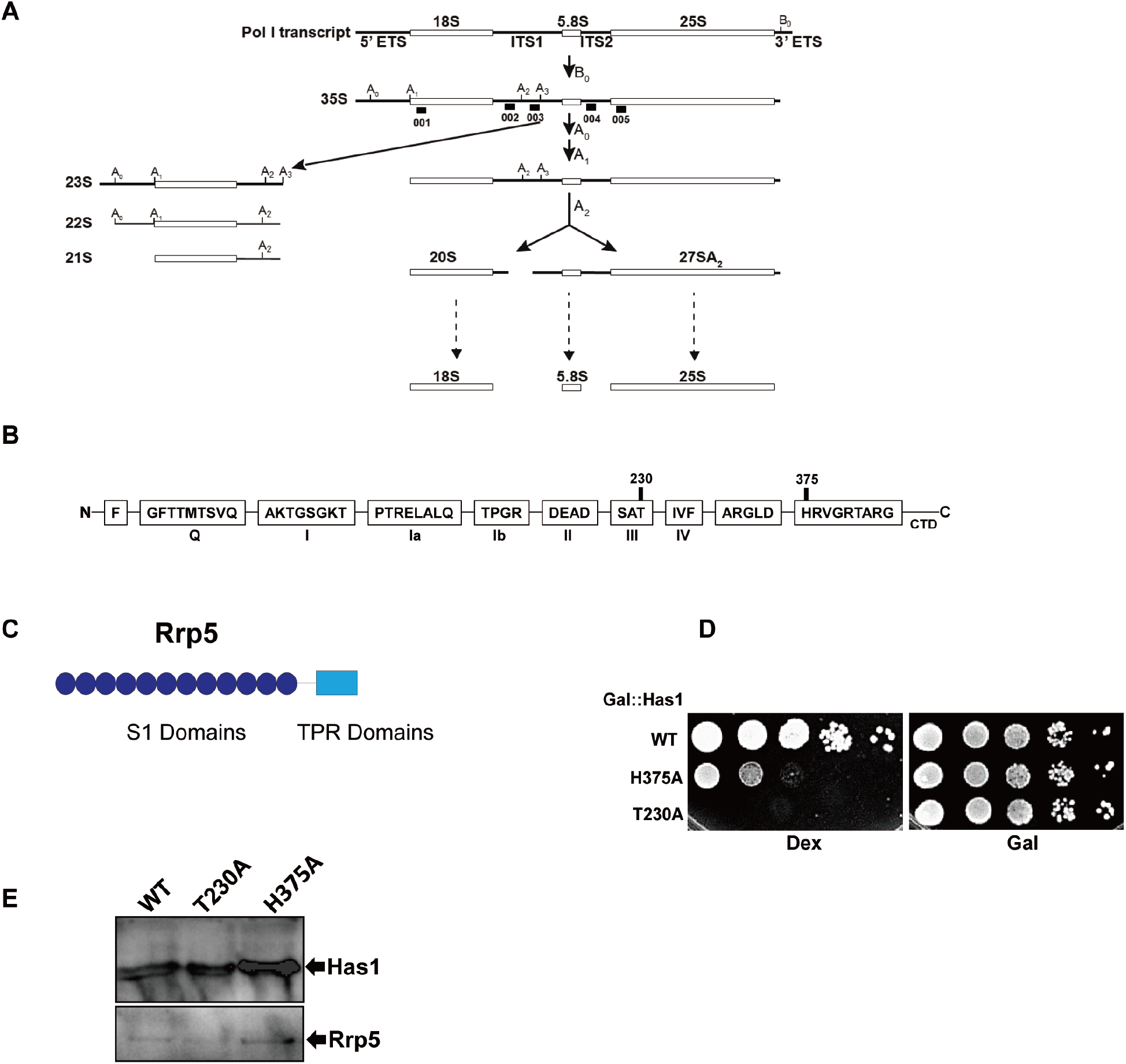
Has1 active site mutants show growth defects. A. rRNA processing scheme and intermediates. The northern probes used herein are indicated with a bar. B. A scheme of the conserved sequence motifs in DEAD-box ATPases shows the location of Has1 active site mutants. C. Schematic representation of the Rrp5 domain organization. D. Serial dilution growth assay of yeast containing Has1 under galactose-inducible control promoter and supplemented with the indicated plasmids. E. Western blot analysis of Has1 expression of total cell protein from Gal:Has1 cells expressing plasmid-encoded Has1 WT or mutants. Rrp5 serves as loading control.

**Figure S2.**
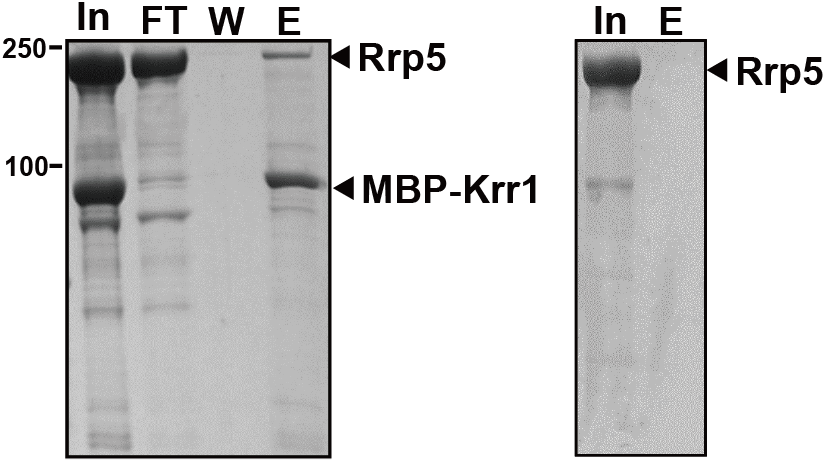
Rrp5 binds Rrp36 and Krr1 in distinct regions of the assembling subunit. Left: Recombinant MBP-Krr1 immobilized on amylose resin interacts with recombinant purified Rrp5_FL. Right: Rrp5 dose not bind amylose resin in the absence of MBP-RrpKrr1. In: Input; FT: Flow through; W: Wash; E: Elution.

**Figure S3.**
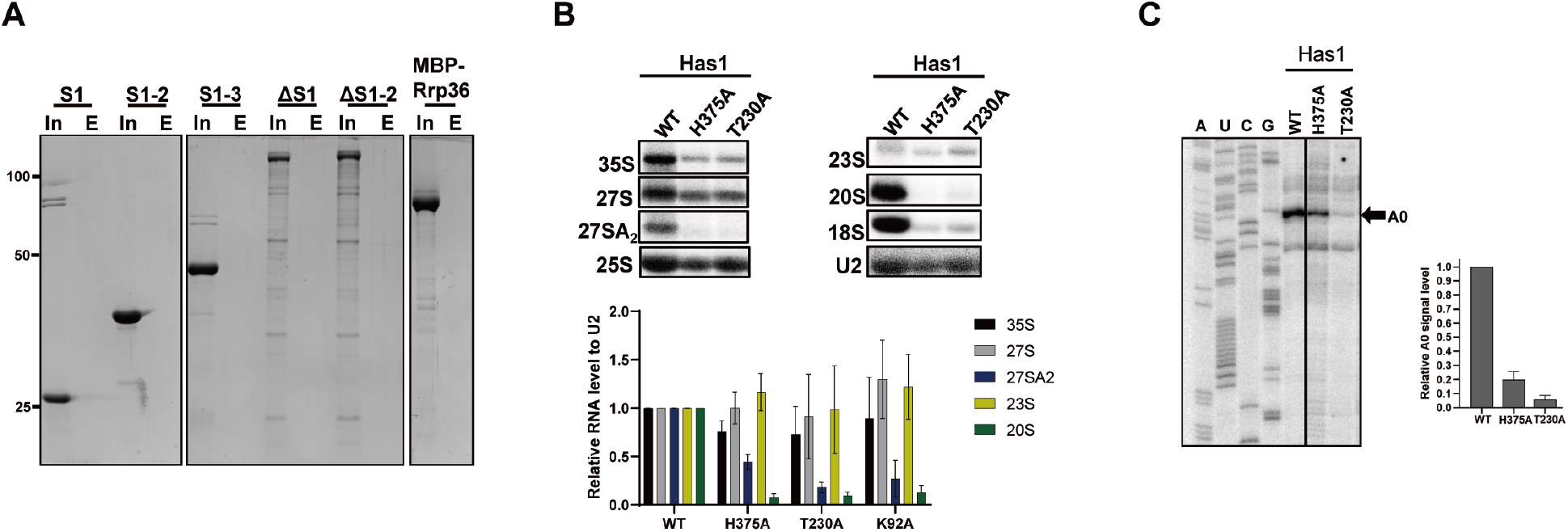
Inactivation of Has1 blocks the first rRNA processing step at site A_0_. A. Left and middle: Rrp5 fragments from Figure 2 do not bind amylose resin in the absence of MBP-Rrp36. Right: MBP-Rrp36 from Figure 2 do not bind Ni-NTA resin in the absence of Rrp5_FL. B. Northern blot analysis of rRNA processing intermediates from Has1-depleted cells expressing plasmid-encoded wild type (WT) Has1 or active site mutants. U2 serves as loading control. Quantification of Northern blots is shown below the blots. rRNA levels were normalized to U2 levels and to Has1 WT. The data are averages from 3 replicate experiments and error bars show standard deviation of the mean. C. Analysis of A_0_ cleavage levels by reverse transcription. RNA was extracted from the Utp10-TAP pull-down elution of Has1 WT or mutants. Shown below is the quantification of A_0_ cleavage levels, which was normalized to the full extension band (not shown). The data are averages from 2 biological replicate experiments and error bars show standard deviation of the mean.

**Figure S4.**
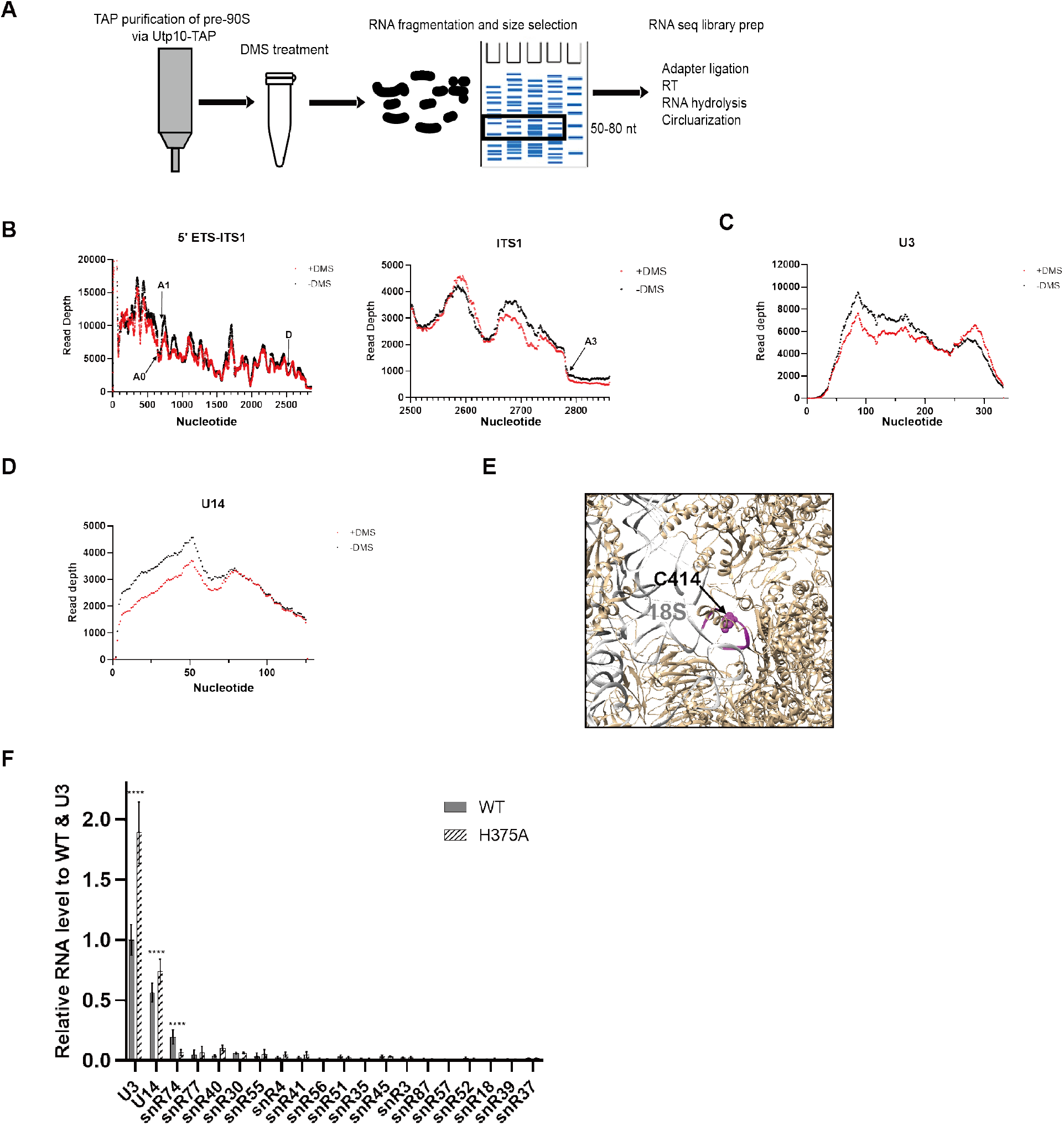
DMS MaPSeq analysis of Has1_H375A. A. A scheme shows sample preparation for DMS-MaPseq. B. DMS-MaPseq read depth of pre-rRNA 5’ETS-ITS1 region (left) and more detail in TS1 region (right). Cleavage site of A0, A1, D and A3 were indicated by black arrows. C. DMS-MaPseq read depth of U3 snoRNA. D. DMS-MaPseq read depth of U14 snoRNA. E. Total reads of snoRNAs were first normalized to the length, then further normalized to U3 in WT. The data are averages from 8 replicate experiments. F. Analysis of A28 modification levels by reverse transcription. Total RNA from Has1 and Rrp5-depleted cells expressing plasmid-encoded indicated was used as template. Shown below is the quantification of A28 signal levels, which was normalized to the full extension band (not shown).

